# Glutamine synthetase mRNA releases sRNA from its 3’UTR to regulate carbon/nitrogen metabolic balance

**DOI:** 10.1101/2022.07.25.501400

**Authors:** Masatoshi Miyakoshi, Teppei Morita, Asaki Kobayashi, Anna Berger, Hiroki Takahashi, Yasuhiro Gotoh, Tetsuya Hayashi, Kan Tanaka

## Abstract

Glutamine synthetase is the key enzyme of nitrogen assimilation, which is encoded in the first cistron of *glnALG* operon and is induced under nitrogen limiting conditions through transcriptional activation by NtrBC in *Salmonella* and *E. coli*. 2-oxoglutarate serves as the carbon skeleton of glutamate and glutamine, but how 2-oxoglutarate fluctuation is controlled in response to nitrogen availability remained unknown. We show that the *glnA* mRNA produces an Hfq-dependent GlnZ sRNA from its 3’
sUTR through RNase E-mediated cleavage. Through a base-pairing mechanism, GlnZ primarily regulates *sucA*, encoding the E1o component of 2-oxoglutarate dehydrogenase. In the cells grown on glutamine as the nitrogen source, the endogenous GlnZ represses the expression of SucA to redirect the carbon flow from the TCA cycle to the nitrogen assimilation pathway. This study also clarifies that the release of GlnZ sRNA from the *glnA* mRNA by RNase E is essential for the post-transcriptional regulation of *sucA*, and thus the mRNA coordinates the two independent functions to balance the supply and demand of the fundamental metabolites.

## INTRODUCTION

Nitrogen is an essential element of the cell. Ammonia is the energetically most preferable nitrogen source and is assimilated via either glutamate dehydrogenase (GDH) or glutamine synthetase (GS) and glutamine 2-oxoglutarate synthetase (GOGAT) pathway (Reitzer, 2003). GDH catalyzes the reductive amination of 2-oxoglutarate (2-OG, a.k.a. α-ketoglutarate) to glutamate (Glu) without ATP consumption. In contrast, GS catalyzes the amidation of Glu to form glutamine (Gln) using one molecule of ATP, and then GOGAT transfers the amide group reductively to 2-OG to generate two Glu molecules, yielding one net Glu from 2-OG (Fig. 1A). Nitrogen is limiting for most bacteria in freshwater, marine, and terrestrial ecosystems (Elser *et al*, 2007) and mammalian large intestines (Reese *et al*, 2018). For a facultative intracellular pathogen *Salmonella enterica, glnA* is essential during invasion and proliferation in the host cells (Popp *et al*, 2015; Klose & Mekalanos, 1997; Aurass *et al*, 2018).

**Fig. 1.**
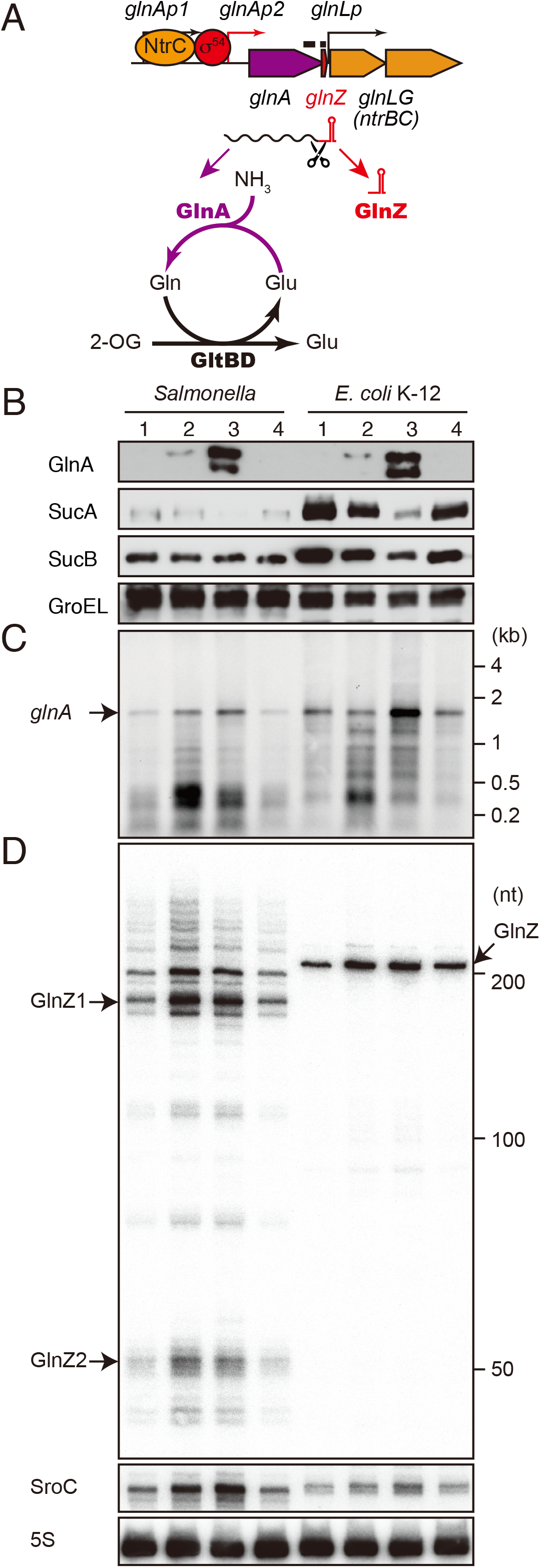
*glnA* mRNA expresses both GlnA protein and GlnZ sRNA. (A) The genetic structure of *glnALG* operon and nitrogen assimilation pathway. The transcription of *glnA* mRNA from σ^54^-dependent promoter *glnAp2* is activated by the NtrB/NtrC two-component system, which is encoded downstream of *glnA*. GlnA (GS) catalyzes the amidation of Glu to form glutamine (Gln) using one molecule of ATP, and GltBD (GOGAT) transfers the amide group of Gln to 2-OG to generate two Glu molecules. GlnZ sRNA is processed from the 3’sUTR of *glnA* mRNA. The black bars above the genes indicate the location of probes used for northern blots. (B) Expression profiles of OGDH subunits and GS in *S. enterica* and *E. coli* during growth on different nitrogen sources. *S*. Typhimurium SL1344 and *E. coli* BW25113 were grown to exponential phase (OD_600_ ∼0.5) in MOPS media containing 0.2% glucose as the carbon source and the following nitrogen sources; 1, 0.1% ammonium, 2, 0.01% ammonium, 3, 5 mM Gln, 4, 5 mM Gln plus 0.01% ammonium. SucA and SucB were detected by antibodies raised against purified *E. coli* proteins. GlnA was detected by an antibody raised against a synthetic peptide. GroEL served as a loading control. (C and D) Expression profiles of *glnA* mRNA and GlnZ sRNA. The *glnA* mRNA was detected by an equimolar mixture of two RNA probes specific for *Salmonella* and *E. coli*. GlnZ was detected by a common oligonucleotide probe MMO-0416. SroC sRNA was detected by an equimolar mixture of oligonucleotide probes JVO-2907 and JVO-5622. 5S rRNA was detected by MMO-1056 served as a loading control.

To deal with the environmental availability of nitrogen, enterobacteria such as *S. enterica* and *E. coli* have developed a complex regulatory network (van Heeswijk *et al*, 2013). The conserved *glnALG* operon encodes GS along with the NtrB/NtrC (GlnL/GlnG) two-component regulator system (Fig. 1A). The response regulator NtrC is phosphorylated by its cognate sensor kinase NtrB at a low nitrogen state and activates RpoN (σ^54^)-dependent promoters by binding at specific enhancer-like sequences (Ninfa & Magasanik, 1986; Ninfa *et al*, 1987). The *glnALG* operon contains three promoters, *glnAp1, glnAp2*, and *glnLp*, and the proximal σ^54^-dependent promoter *glnAp2* is activated by NtrC when phosphorylated by NtrB in nitrogen-poor conditions while the other σ^70^-dependent promoters are repressed by NtrC (Ueno-Nishio *et al*, 1984; Reitzer & Magasanik, 1985). The NtrB/NtrC system can be stimulated when the cells utilize Gln or low concentrations of ammonium as a sole nitrogen source (Schumacher *et al*, 2013), but controversially it has been postulated that a low intracellular Gln level is perceived as the nitrogen limitation to induce the transcription of *glnA* (Ikeda *et al*, 1996).

2-OG serves as both the TCA cycle intermediary metabolite and the carbon skeleton of Glu and Gln and thus has been proposed as a plausible signaling compound coordinating carbon and nitrogen metabolism (Huergo & Dixon, 2015). 2-OG accumulates rapidly upon nitrogen limitation from 0.2 to 10 mM (Yuan *et al*, 2009) and directly blocks glucose uptake by inhibiting enzyme I of the phosphoenolpyruvate phosphotransferase system (PTS) (Doucette *et al*, 2011). 2-OG also inhibits cAMP synthesis by adenylate cyclase (You *et al*, 2013). However, it is currently unknown how the cells adjust the 2-OG levels in response to nitrogen availability.

The genes under the direct control of RpoN and NtrC have been identified in *S. enterica* serovar Typhimurium (Samuels *et al*, 2013; Bono *et al*, 2017) and *E. coli* K-12 strains (Brown *et al*, 2014; Zimmer *et al*, 2000), but none of the genes are implicated in the central carbon metabolism. In general, alternative sigma factors drive the transcription of small RNAs, which regulate the expression of target genes *in trans* at the post-transcriptional level, to extend the regulatory networks of various stress responses (Beisel & Storz, 2010; Mandin & Guillier, 2013; Hör *et al*, 2020). However, little is known about RpoN-dependent sRNAs directly involved in nitrogen metabolism (Prasse & Schmitz, 2018). The only RpoN-dependent sRNA known so far in enterobacteria is GlmY, which accumulates upon glucosamine-6-phosphate depletion through transcriptional activation by the GlrK/GlrR two-component system (Urban *et al*, 2007; Reichenbach *et al*, 2009). Since *S. enterica* and *E. coli* contain a dozen two-component systems associated with RpoN (Studholme, 2002; Hartman *et al*, 2016), there may be more than one RpoN-dependent sRNA.

In addition to intergenic regions of bacterial genomes, mRNA 3’
sUTRs have emerged as a reservoir of an emerging class of post-transcriptional regulators (Miyakoshi *et al*, 2015b; Adams & Storz, 2020; Ponath *et al*, 2022). Indeed, the *gltIJKL* operon encoding the Glu/Asp ABC transporter is transcribed from σ^70^- and σ^54^-dependent promoters, and the *gltI* mRNA with a Rho-independent terminator is processed by RNase E to release SroC sRNA from its 3’UTR (Miyakoshi *et al*, 2015a). SroC functions as the sponge RNA of GcvB sRNA, which regulates >50 genes mostly involved in amino acid metabolism and transport (Miyakoshi *et al*, 2022). Likewise, *glnA* contains a Rho-independent terminator with a moderate strength upstream of *glnLp* (Ueno-Nishio *et al*, 1984), and thus the *glnA* mRNA is likely to efficiently bind to RNA chaperone Hfq at its 3’ end (Chen *et al*, 2019). An sRNA has been detected in the 3’UTR of *glnA* mRNA in *E. coli*, renamed as GlnZ (Kawano *et al*, 2005; Walling *et al*, 2022), and also in the same locus of *S. enterica* (Sittka *et al*, 2009). Therefore, GlnZ is regarded as a RpoN-dependent sRNA in charge of the noncoding arm of the NtrC regulon.

Here we show that GlnZ sRNA derived from the 3’
sUTR of *glnA* mRNA represses the expression of *sucA* encoding the E1o component of 2-oxoglutarate dehydrogenase (OGDH) and thus functions as the post-transcriptional regulator of the TCA cycle branch point directly linking carbon and nitrogen metabolism in *S. enterica* and *E. coli*. In parallel, the NtrC-dependent transcriptional regulator Nac, which is absent in *S. enterica*, also represses *sucA* specifically in *E. coli*. Moreover, we demonstrate that RNase E-mediated cleavage of GlnZ from the *glnA* mRNA is a prerequisite for targeting the *sucA* mRNA.

## RESULTS

### Expression profiles of OGDH subunits and the *glnA* products upon nitrogen limitation

As 2-OG accumulates upon nitrogen limitation (Yuan *et al*, 2009), we hypothesized that the expression level of OGDH is reduced to cause a bottleneck in the TCA cycle. To this end, we analyzed the expression levels of OGDH subunits, SucA and SucB, in *S*. Typhimurium SL1344 and *E. coli* K-12 by western blot. Throughout this study, we used 0.2% glucose MOPS minimal medium whose nitrogen source was substituted by either 0.1% ammonium, 0.01% ammonium, 5 mM Gln, or 5 mM Gln plus 0.01% ammonium. In line with the previous report (Schumacher *et al*, 2013), GS was induced with the lower concentration of ammonium and much further during growth on Gln both in *S. enterica* and *E. coli* (Fig. 1B). Addition of ammonium into the Gln medium hindered the induction, i.e. nitrogen catabolite repression. In contrast, the expression level of SucA was strikingly reduced during growth on Gln and counter-correlated with that of GS in both strains. In *E. coli*, SucB exhibited a similar expression pattern to SucA. However, SucB was expressed constantly during growth under the four growth conditions in *S. enterica*, although *sucA* and *sucB* are cotranscribed. This result suggests that the expression of *sucA* is regulated at the post-transcriptional level in response to nitrogen availability.

To clarify whether *glnA* mRNA and GlnZ sRNA are also induced upon nitrogen limitation, total RNAs were analyzed by northern blotting. As expected, the *glnA* mRNA was induced during growth on Gln as the sole nitrogen source in *S. enterica* and *E. coli* (Fig. 1C). At the low concentration of ammonium, degradation products in the 3’
s region of *glnA* CDS accumulated as the cells starve for ammonium and stop growth. In the sRNA fractions, we detected several GlnZ isoforms in the *S. enterica* total RNA (Fig. 1D). The size of the most abundant transcript matches that of the *glnA* 3’UTR spanning 185 nucleotides (nt), given the transcription stops at the Rho-independent terminator with 6 U residues. We also detected ∼50 nt short transcripts, which are likely produced by RNase E-mediated cleavage (Chao *et al*, 2017). Here the most abundant transcript is designated as GlnZ1 and the shorter transcript as GlnZ2. In contrast, a ∼200-nt single transcript was seen in the *E. coli* K-12 total RNA using the same probe, which hybridizes the 5’ end region of K-12 GlnZ. In both *S. enterica* and *E. coli* K-12, the GlnZ transcripts were abundant under the two nitrogen limiting conditions. Concomitantly, we observed similar induction of *S. enterica* SroC in the same samples (Fig. 1D). In contrast, the induction of SroC was only marginal in *E. coli* K-12 due to the IS*5* insertion upstream of *gltI* (Zinser *et al*, 2003).

### Sequence comparison of the *glnA-glnLG* intergenic region among *Enterobacteriaceae*

As the fragment patterns of GlnZ transcripts were different between *S. enterica* and *E. coli*, we compared the sequence of *glnA* 3’
sUTRs among *Enterobacteriaceae*. The 185-nt *glnA* 3’UTR is >99% identical among *Salmonella* species/subspecies except for *S. enterica* subsp. *diarizonae* and *salamae* (Fig. 2A). In contrast, *E. coli glnA* 3’UTRs are phylogenetically diverse and can be classified into at least three types. *E. coli* K-12 and several pathogenic strains harbor the 197-nt *glnA* 3’UTR (K-12 type). Many *E. coli* pathogenic strains represented by O157 carry a shorter *glnA* 3’UTR of 85 nt (O157 type), while the others represented by O111 have a longer 3’UTR of 230 nt (O111 type), the latter of which has acquired a 98-nt insertion sequence flanked by 5’-AGGUUCAAA-3’ direct repeats into the O157-type *glnA* 3’UTR (Fig. 2B). The *glnA* 3’UTRs of the other species in the genus *Escherichia, E. albertii* and *E. fergusonii*, are almost identical to the K-12 and O157 types, respectively.

**Fig. 2.**
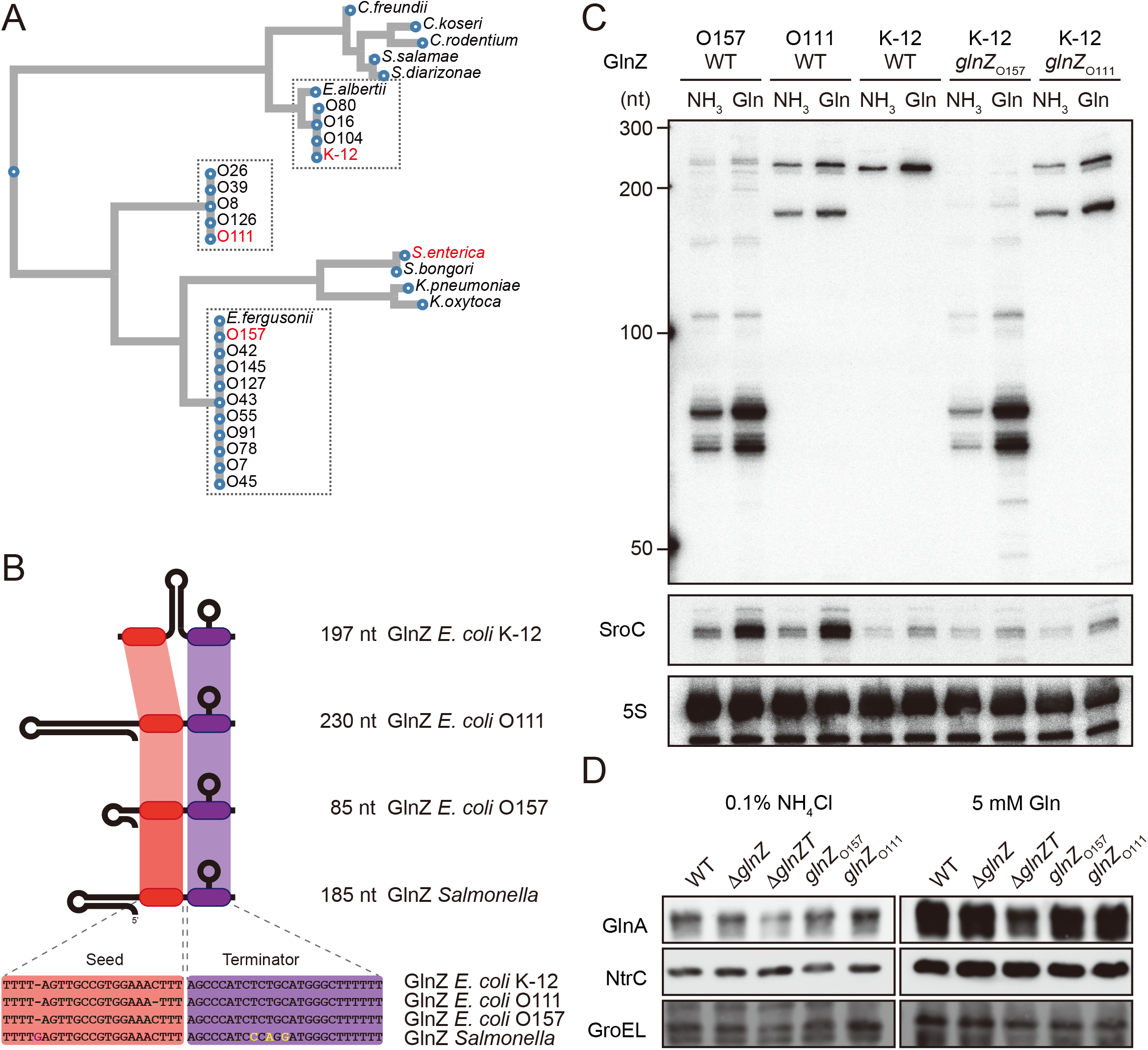
Variation of GlnZ sRNAs among *Enterobacteriaceae*. (A) Multiple sequence alignment of *glnA* 3’sUTRs using CLUSTALW program (https://www.genome.jp/tools-bin/clustalw). (B) The location of conserved seed region and Rho-independent terminator in different types of GlnZ sRNAs. The extra G nucleotide and variable nucleotides found in the terminator of *Salmonella* GlnZ are indicated in purple and yellow letters. (C) *E. coli* strains express different types of GlnZ sRNAs. *E. coli* O157, O111, and K-12 strains were grown to exponential phase (OD600 ∼0.5) in MOPS media containing 0.2% glucose as the carbon source and either 0.1% ammonium or 5 mM Gln as the nitrogen source. GlnZ and SroC sRNAs were detected by MMO-0416 and JVO-5622, respectively. 5S rRNA detected by MMO-1056 served as a loading control. (D) The difference in 3’UTR sequence does not affect the expression of downstream NtrC. *E. coli* K-12 strains, WT, Δ*glnZ*, Δ*glnZT, glnZ*_O157_, and *glnZ*_O111_, were grown to exponential phase (OD600 ∼0.5) in MOPS media containing 0.2% glucose as the carbon source and either 0.1% ammonium or 5 mM Gln as the nitrogen source. GlnA was detected by an antibody raised against a synthetic peptide. NtrC was chromosomally tagged with 3xFLAG and detected by α-FLAG antibody. GroEL served as a loading control.

To verify whether these strains generate different types of GlnZ transcripts, total RNAs extracted from the O157 and O111 cells were analyzed by northern blotting. We detected GlnZ sRNAs corresponding to the *glnA* 3’
sUTRs in size and their processed species, which were induced when Gln was used as the sole nitrogen source (Fig. 2C). This result is consistent with the study by Walling, Kouse et al. (Walling *et al*, 2022) but not the previous report by Jia et al. (Jia *et al*, 2021). In support, SroC was also induced under the nitrogen limiting condition in the two pathogenic *E. coli* strains devoid of IS*5* insertion upstream of *gltI*.

The structures of *E. coli glnA* 3’
sUTRs might affect the expression of the downstream *glnLG* genes encoding NtrBC. In the K-12 strain, the expression of GlnA and NtrC was not affected by the partial deletion of *glnA* 3’UTR retaining the Rho-independent terminator (Δ*glnZ*) (Fig. 2D). In contrast, the level of GlnA was slightly reduced by further deletion of the terminator (Δ*glnZT*) while that of NtrC remained nearly constant. These results suggest that the Rho-independent terminator stabilizes the *glnA* mRNA but not the *glnALG* mRNA or that the *glnLG* mRNA is transcribed independently to maintain the level of NtrC. Next, we replaced the *glnA* 3’UTR in the K-12 chromosome with that of O157 or O111. The recombinant strains expressed the heterologous GlnZs (Fig. 2C), but the difference in the expression levels of GlnA and NtrC between these strains and the parental K-12 strain was negligible (Fig. 2D). These results indicate that the various *glnALG* operon of *E. coli* strains function equivalently.

### Target identification of GlnZ sRNA in *Salmonella* and *E. coli*

Enterobacterial GlnZ sRNAs contain a conserved sequence, 5’
s-UUGCCGUGGAAA-3’, which is located adjacent to the Rho-independent terminator in *S. enterica* or just downstream of *glnA* stop codon in *E. coli* K-12. Thus, this sequence was assumed to function as the seed region to interact with target mRNAs (Storz *et al*, 2011; Gorski *et al*, 2017). We searched for the complementary sequences to the putative seed region in several enterobacterial genomes using the CopraRNA program (Wright *et al*, 2013). This prediction suggested that the primary target of GlnZ is most likely the *sucA* gene, which encodes the E1o subunit of 2-oxoglutarate dehydrogenase (OGDH). *sucA* is located in the *sdhCDAB-sucABCD* operon, which is regulated by multiple sRNAs but also produces an sRNA regulator SdhX from its 3’UTR (Miyakoshi *et al*, 2019; de Mets *et al*, 2019). The *sdhB-sucA* intergenic region spans 523 nt in *S*. Typhimurium SL1344 and 300 nt in *E. coli* K-12. The sequence complementary to the GlnZ seed region is located upstream of the translation initiation region of *sucA* in *E. coli* and far upstream in *S. enterica* (Fig. 3AB).

**Fig. 3.**
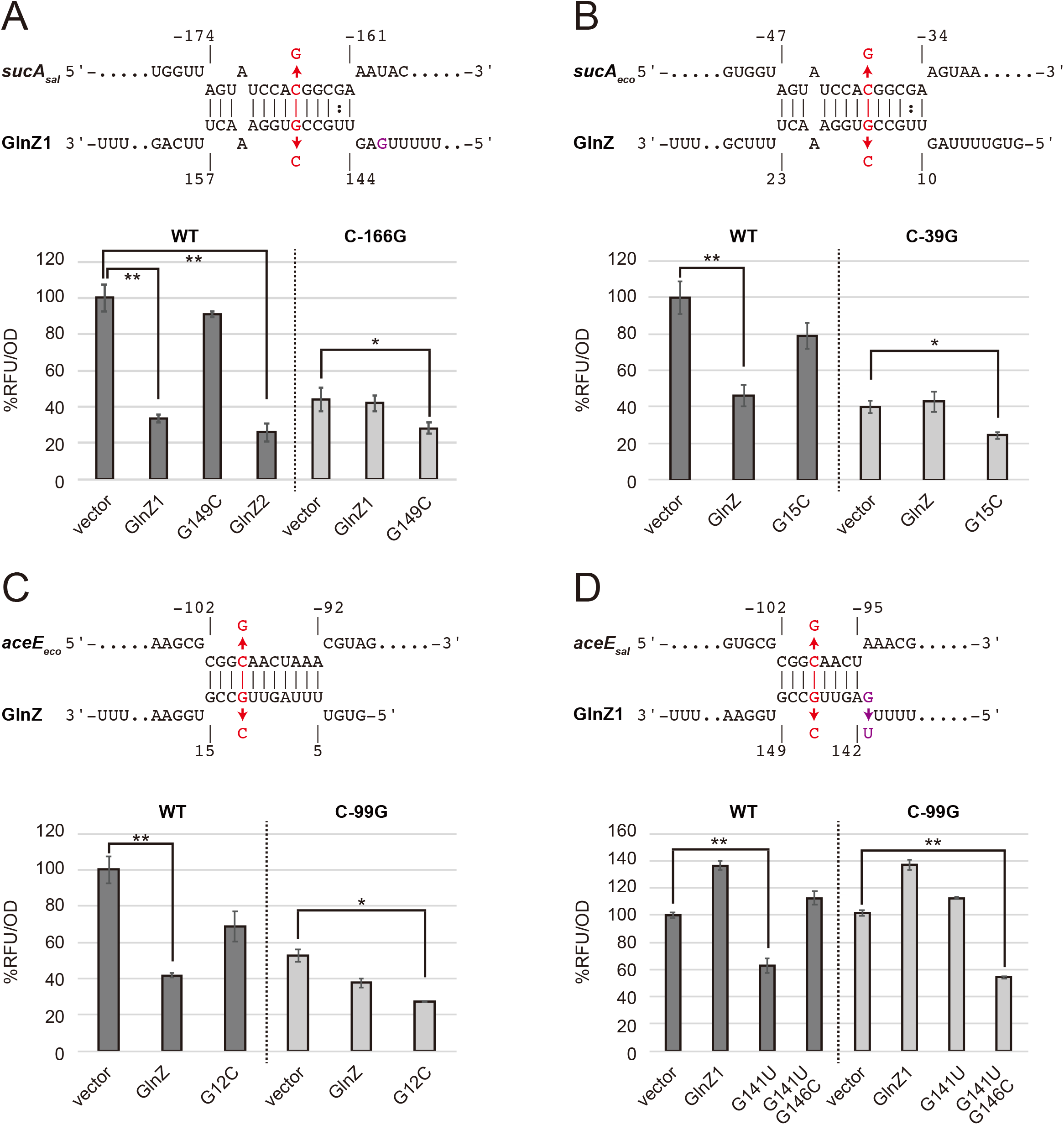
Post-transcriptional regulation of *sucA* (A, B) and *aceE* (C, D). Predicted interactions of GlnZ with the target mRNAs are shown above the bar graphs. The nucleotide numbers relative to the start codon of the target mRNA and the stop codon of *glnA* are shown above and below the nucleotide sequences, respectively. The mutated nucleotides are indicated in red, and the extra G nucleotide found in *Salmonella* GlnZ1 is shown in purple. *E. coli* Δ*glnZ* strain was transformed by GFP translational fusion plasmids along with pJV300 control vector or GlnZ expression plasmids (Supplementary Tables S5-7). Mean relative fluorescence units (RFU) normalized by OD_600_ calculated from biological replicates (n>3) are presented with standard deviation in percentage relative to the vector control. Statistical significance was calculated using one-way ANOVA and denoted as follows: ** *p* < 0.005, * *p* < 0.05.

To verify the post-transcriptional regulation by GlnZ, the *sdhB-sucA* intergenic region was cloned into the pXG30-sf translational fusion vector (Corcoran *et al*, 2012). When either GlnZ1 or GlnZ2 was ectopically expressed from a constitutive promoter, the fluorescence of *S. enterica* SucA::GFP fusion was significantly reduced (Fig. 3A). This result indicates that SucA is repressed by GlnZ1, and the shorter transcript GlnZ2 is sufficient to repress *sucA*. The repression was abrogated when the 149th nucleotide downstream of the *glnA* stop codon was mutated from G to C in GlnZ1 (G149C) and when the 166th nucleotide upstream of the *sucA* start codon was mutated from C to G in *sucA* (C-166G). The repression was restored when GlnZ1 and its target were simultaneously mutated in the complementary nucleotides, demonstrating the post-transcriptional regulation through the base-pairing mechanism. Similarly, *E. coli* GlnZ repressed *sucA* irrespective of the position of seed region in the sRNA and that of target region in the mRNA (Fig. 3B).

To search for additional targets of GlnZ, we performed RNA-seq analysis upon GlnZ1 pulse-expression in *S. enterica* cells exponentially grown in LB medium. Among the 12 genes that showed more than 4-fold downregulation (Supplementary Table S1), *glnP* and *glnQ* constitute the *glnHPQ* operon encoding the Gln ABC transporter. We also found *deoD* encoding a purine nucleoside phosphorylase as a candidate GlnZ target. Using IntaRNA program (Mann *et al*, 2017), both *glnP* and *deoD* mRNAs were predicted to base-pair with the GlnZ seed region. Using the pXG30-sf constructs of *glnHP* and *deoBD* intergenic regions, we verified that the expression of *glnP* and *deoD* was repressed by *S. enterica* GlnZ1 and GlnZ2 through the base-pairing mechanism (Supplementary Figure S1). It is noteworthy that the *glnP* target region is conserved in *E. coli* but that of *deoD* is not.

Next, we looked into the RNA-RNA interactome datasets in *E. coli* K-12 and UPEC O127:H6 (Melamed *et al*, 2016, 2020; Pearl Mizrahi *et al*, 2021). In the cells grown in LB medium, the majority of GlnZ transcripts interacted with the *sdhB-sucA* intergenic region. Interestingly, GlnZ was also found to associate with the *pdhR-aceE* intergenic region. Using the translational fusion of *pdhR-aceE*, we observed significant repression of AceE::GFP fusion by *E. coli* GlnZ (Fig. 3C). The repression was relieved by the G12C mutation in GlnZ, but was restored by its complementary C-99G mutation in *aceE*, showing that GlnZ regulates *aceE* through base-pairing far upstream of the translation initiation region. However, the expression of *aceE* was not repressed by *Salmonella* GlnZ1 (Fig. 3D). We reasoned that the seed region of GlnZ1 contains the insertion of G at the 143rd nucleotide, and thus the interaction with *aceE* is weaker than the *E. coli* GlnZ. When the nucleotide is substituted by U (G143U), GlnZ1 gained the inhibitory effect on *S. enterica aceE* by extending the hybrid by 3 bp (Fig. 3D). Overall, GlnZ post-transcriptionally regulates *sucA* and *glnP* in common, but a mutation adjacent to the seed region brings about a species-specific regulation.

### GlnZ represses *sucA* during growth on Gln as the sole nitrogen source

As the post-transcriptional regulation by GlnZ through the base-pairing mechanism has been verified, we evaluated the effect of endogenous GlnZ on its primary target *sucA*. To this end, we constructed mutant strains partially lacking the *glnA* 3’
sUTRs (Δ*glnZ*) of *S*. Typhimurium SL1344 and *E. coli* K-12 and analyzed the SucA level by western blotting. These deletion mutants grew in the minimal media similarly to the respective WT strains. During growth on Gln, the *S. enterica* Δ*glnZ* strain expressed >1.7-fold higher levels of SucA than WT during growth on Gln (Fig. 4A), indicating that GlnZ repressed SucA during growth on the non-preferable nitrogen source. Likewise, we observed a modest increase in the SucA expression level by deleting *glnZ* in *E. coli* during growth on Gln as the sole nitrogen source (Fig. 4B).

**Fig. 4.**
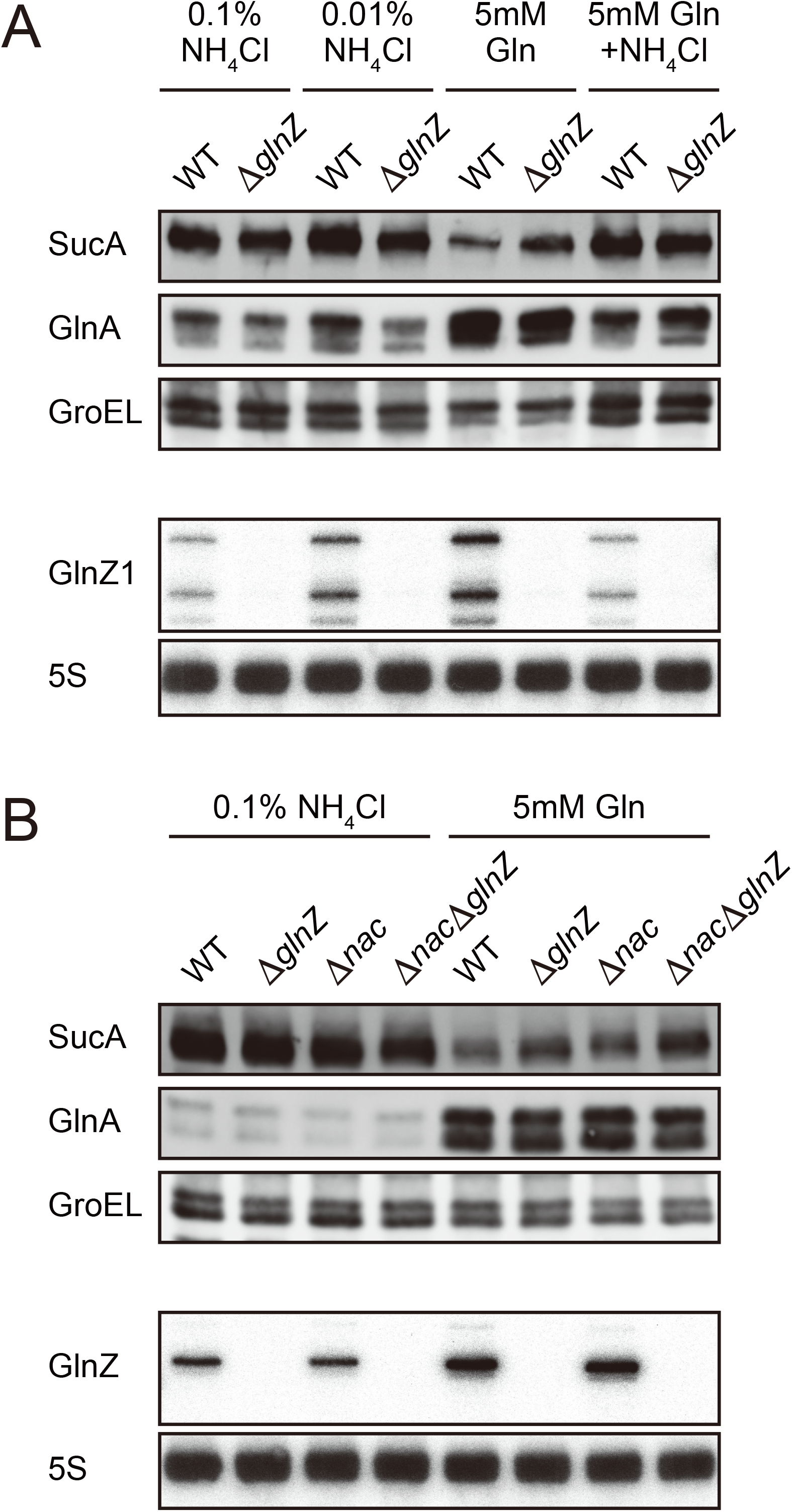
Repression of SucA by endogenous GlnZ. (A) GlnZ post-transcriptionally represses the expression OGDH subunits in *Salmonella* during growth on Gln as the nitrogen source. *Salmonella* WT and Δ*glnZ* strains were grown to exponential phase (OD_600_ ∼0.5) in MOPS minimal medium containing 0.2% glucose as the carbon source and different nitrogen sources; 0.1% ammonium, 0.01% ammonium, 5 mM Gln, and 5 mM Gln plus 0.01% ammonium. (B) GlnZ and Nac independently repress the expression OGDH subunits in *E. coli* during growth on Gln as the nitrogen source. *E. coli* WT, Δ*glnZ*, Δ*nac*, and Δ*glnZ*Δ*nac* strains were grown to exponential phase (OD_600_ ∼0.5) in MOPS minimal medium containing 0.2% glucose as the carbon source and either 0.1% ammonium or 5 mM Gln as the nitrogen source. The expression of OGDH subunits, SucA and SucB, and GS was analyzed by western blots (upper panels), and that of GlnZ sRNA was analyzed by northern blots (bottom panels). GroEL served as a loading control for western blots; 5S rRNA for northern blots.

Still, the difference was much more significant when compared between the poor and rich nitrogen sources, implicating additional regulators in the control of SucA expression. Recently, the global regulator of nitrogen assimilation control (Nac), which is absent in *S. enterica* (Muse & Bender, 1998), was found to bind the *sucA* promoter in *E. coli* (Aquino *et al*, 2017). In line with this finding, SucA and SucB exhibited a similar expression pattern during growth with different nitrogen sources in *E. coli*, while the level of SucB was constant in *S. enterica* (Fig. 1B). We verified that the deletion of *nac* alone increased the SucA level without affecting the GlnZ level, and the double deletion of *nac* and *glnZ* further raised the SucA level by >2.5-fold compared to WT (Fig. 4B). As Nac and GlnZ are induced through transcriptional activation by NtrC, *E. coli* utilizes the transcriptional and post-transcriptional regulators to repress the same target in parallel.

### RNase E is indispensable for GlnZ biogenesis

A 3’
sUTR-derived sRNA is either transcribed from an ORF-internal promoter as a primary transcript or released from an mRNA by a ribonuclease (Miyakoshi *et al*, 2015b). To clarify in which pathway GlnZ is generated, we ectopically expressed the *glnA* mRNA of *S. enterica* and *E. coli* from an arabinose-inducible pBAD promoter in the *E. coli* Δ*glnZ* background. With this plasmid expression system, we detected processed fragments as in the endogenous transcripts but virtually no transcripts in the absence of L-arabinose (Fig. 5A), confirming that GlnZ is generated from the *glnA* mRNA. GlnZ was similarly produced from a precursor transcript with the 90-nt extension at 5’ end (GlnZ+90), suggesting that upstream sequence of at least 90 nt is sufficient for GlnZ processing.

**Fig. 5.**
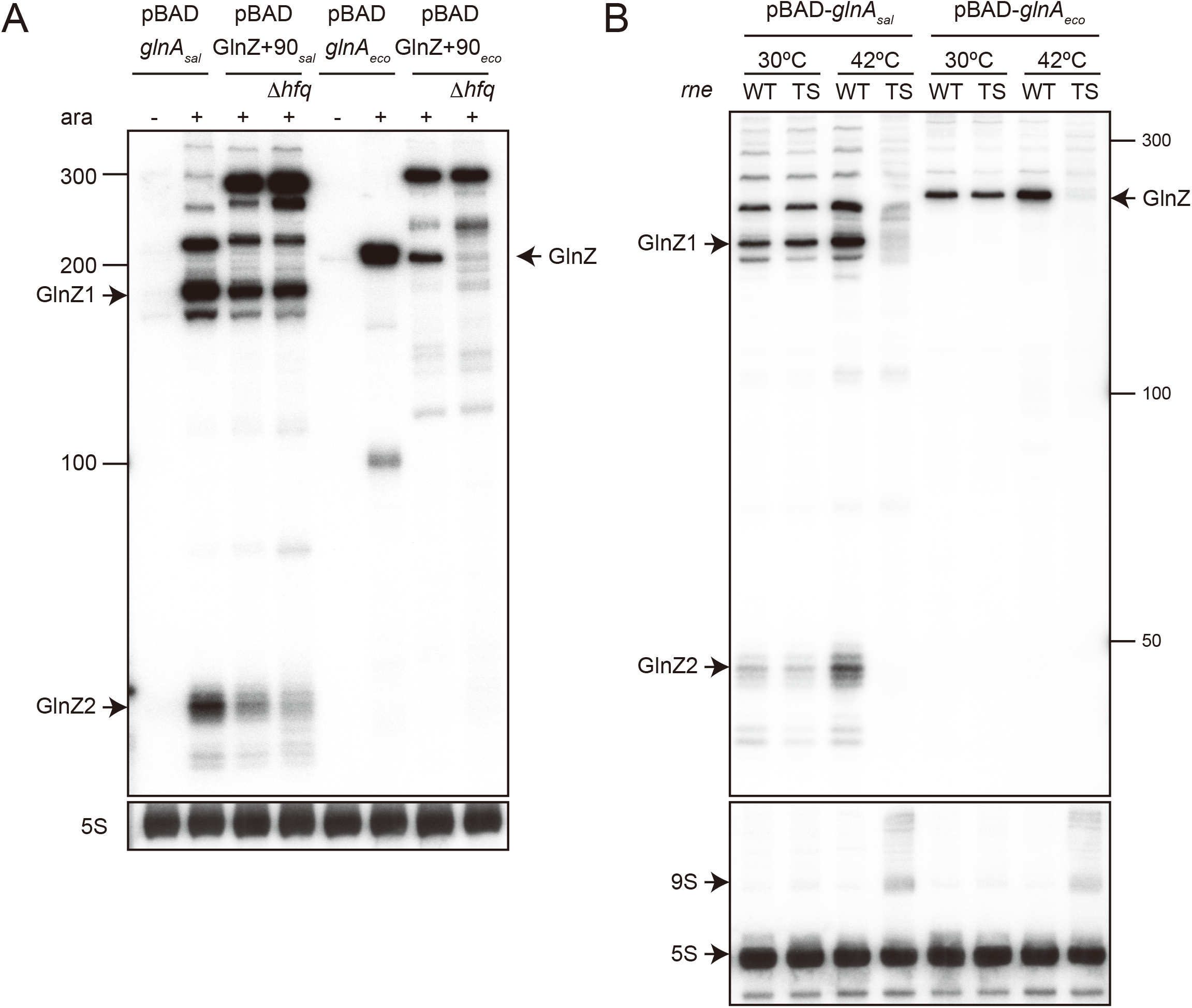
RNase E is essential for GlnZ biogenesis. (A) GlnZ is generated primarily from the *glnA* mRNA. *E. coli* Δ*glnZ* and Δ*glnZ*Δ*hfq* strains harboring pBAD-*glnA*_sal_, pBAD-preGlnZ_sal_, pBAD-*glnA*_eco_, or pBAD-preGlnZ_eco_ were grown to OD600 ∼0.5 at 37ºC and further incubated for 10 min in the presence (+) or absence (−) of 0.2% L-arabinose. (B) RNase E is essential for the processing of *glnA* mRNA. *E. coli* Δ*glnZ* (WT) and Δ*glnZ ams*-1 (TS) strains harboring pBAD-*glnA*_sal_ or pBAD-*glnA*_eco_ were grown to OD600 ∼0.5 at 30ºC and split into two flasks. The flasks were incubated at either 30ºC or 42ºC for 30 min and further incubated for 10 min after adding 0.2% L-arabinose. The size is estimated by DynaMarker RNA Low II ssRNA fragment.

Since *E. coli* GlnZ is classified as a class I sRNA which binds to proximal and rim faces of RNA chaperone Hfq (Kavita *et al*, 2022), Hfq is implicated in the stability of GlnZ. However, the levels of ectopically expressed GlnZ+90 were not altered (Fig. 5A). In the *hfq* mutant, the abundance of *Salmonella* GlnZ1 was not significantly altered, while that of GlnZ2 was slightly reduced. In contrast, no *E. coli* GlnZ was produced from the precursor in the *hfq* mutant despite the comparable expression levels of GlnZ+90 (Fig. 5A). This result suggests that the processing of *E. coli* GlnZ, but not its stability, strictly depends on Hfq.

Several RNase E cleavage sites have been identified in the *glnA* 3’
sUTR in *S. enterica* (Chao *et al*, 2017). To clarify the requirement of RNase E, we pulse-expressed the *glnA* mRNA of *S. enterica* and *E. coli* for 10 min either in the *E. coli* WT or temperature-sensitive RNase E mutant *ams-1* (TS). The fragment patterns were comparable between WT and TS strains at the permissible temperature. At the non-permissible temperature, the TS strain exhibited virtually no accumulation of processed GlnZ sRNAs (Fig. 5B). Thus, we conclude that RNase E is required for GlnZ biogenesis from the *glnA* mRNAs.

### sRNA release from its parental mRNA is essential for target regulation

GlnZ sRNA is crucial for the post-transcriptional regulation of *sucA*. Still, the parental *glnA* mRNA can also be the regulator because it contains both the seed region and Rho-independent terminator required for Hfq-mediated interaction. To this end, we analyzed whether GlnZ processing is a prerequisite for its target regulation using the *E. coli* K-12 pBAD-*glnA* variants. We identified two RNase E cleavage sites 1 and 2, located downstream and upstream of the *glnA* stop codon, which are conformable to the consensus motif (Chao *et al*, 2017), and introduced mutations into the pBAD-*glnA*_eco_ plasmid to disrupt the processing by RNase E without affecting the seed region (Fig. 6A). The mutation at site 1 (mut1) strongly inhibited the processing but slightly stimulated the cleavage at site 2 compared to WT while the mutation at site 2 (mut2) did not affect the cleavage at site 1 (Fig. 6B). Simultaneous mutations at the two cleavage sites (mut3) almost completely abrogated the processing of the *glnA* mRNA (Fig. 6B). This result suggests that RNase E processively cleaves the two sites of the *glnA* mRNA in the 5’
s to 3’ direction.

**Fig. 6.**
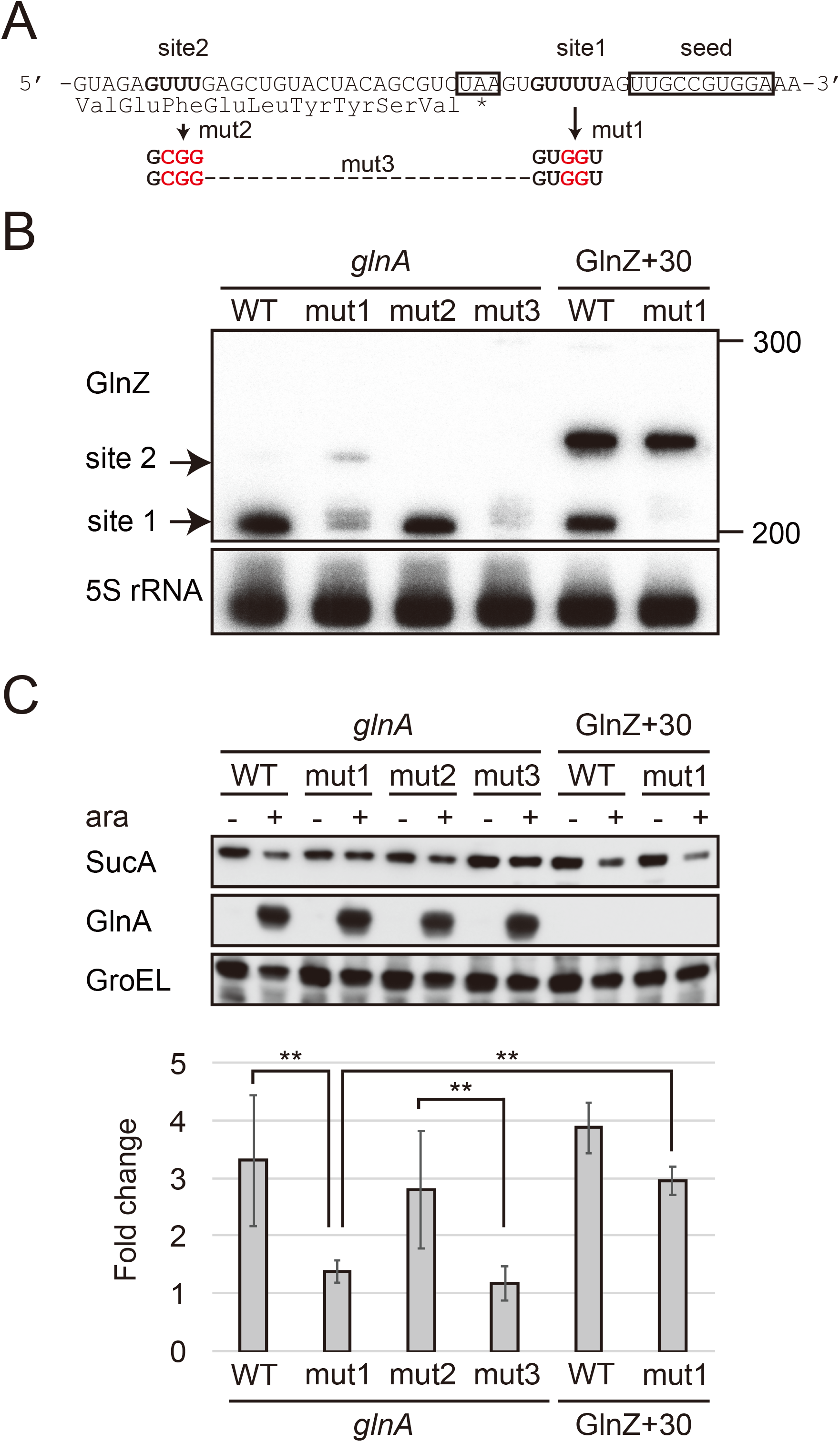
GlnZ release from the *glnA* mRNA is necessary for target repression. (A) The nucleotide sequence of RNase E cleavage sites in *E. coli* K-12 *glnA* mRNA. The C-terminal amino acid sequence of GlnA is shown below the nucleotide sequence. The *glnA* stop codon and the GlnZ seed region are boxed. The mutated nucleotides are indicated in red. (B) The processing of GlnZ is abrogated by the mutations in RNase E cleavage sites. GlnZ sRNA processed from either the *glnA* mRNA or the GlnZ_227_ precursor sRNA was analyzed by northern blot. 5S rRNA served as a loading control. (C) The processing of GlnZ is required for the repression of SucA in the form of mRNA but not the precursor sRNA. Expression levels of SucA and GlnA were analyzed by western blot. GroEL served as a loading control. *E. coli* Δ*glnZ* strains harboring pBAD expression plasmids were grown to exponential phase (OD_600_ ∼1.0) in LB medium in the absence (-) or presence of 0.01% L-arabinose (+). Bar graph represents the fold change of SucA repression by the *glnA* mRNA induced by arabinose calculated from biological replicates (n>5) with standard deviation. Statistical significance was calculated using one-way ANOVA and denoted as **, *p* < 0.005.

Next, we analyzed the protein levels upon ectopic expression of the pBAD-*glnA* variants. Western blot analysis showed that these plasmids expressed GS at strikingly higher levels than the endogenous level (Fig. 6C). This result also confirms that mutations in RNase E cleavage sites did not affect the translation of GlnA. Ectopic expression of the WT *glnA* mRNA strikingly repressed SucA, but when mut1 was introduced into the *glnA* mRNA, SucA was no longer repressed (Fig. 6C). Notably, this mutation also abrogated the processing of precursor GlnZ sRNA with the 30-nt extension at the 5’
s end (GlnZ+30) but did not significantly affect the repression of SucA (Fig. 6BC). This result indicates that GlnZ needs to be separated from its parental mRNA to regulate its target mRNA. If the *glnA* mRNA is not processed properly, the translation might interfere with the base-pairing interaction with the target mRNA.

## DISCUSSION

This study shows that the GlnZ sRNA post-transcriptionally represses the *sucA* mRNA encoding OGDH to redirect 2-OG from the TCA cycle into the GS-GOGAT pathway (Fig. 7). Thus, both the sRNA and protein expressed from the same mRNA act independently to balance the supply and demand of the fundamental intermediates for carbon and nitrogen metabolism. Specifically in *E. coli*, Nac reinforces the repression of *sucA* at the transcriptional level in parallel. Importantly, GlnZ is generated by RNase E-dependent cleavage from the 3’
sUTR of *glnA* mRNA. Although the *glnA* mRNA possesses the regulatory module at its 3’UTR, it is unable to regulate its target mRNA *in trans*. Thus, the sRNA needs to be separated from its parental mRNA to regulate its target mRNA.

**Fig. 7.**
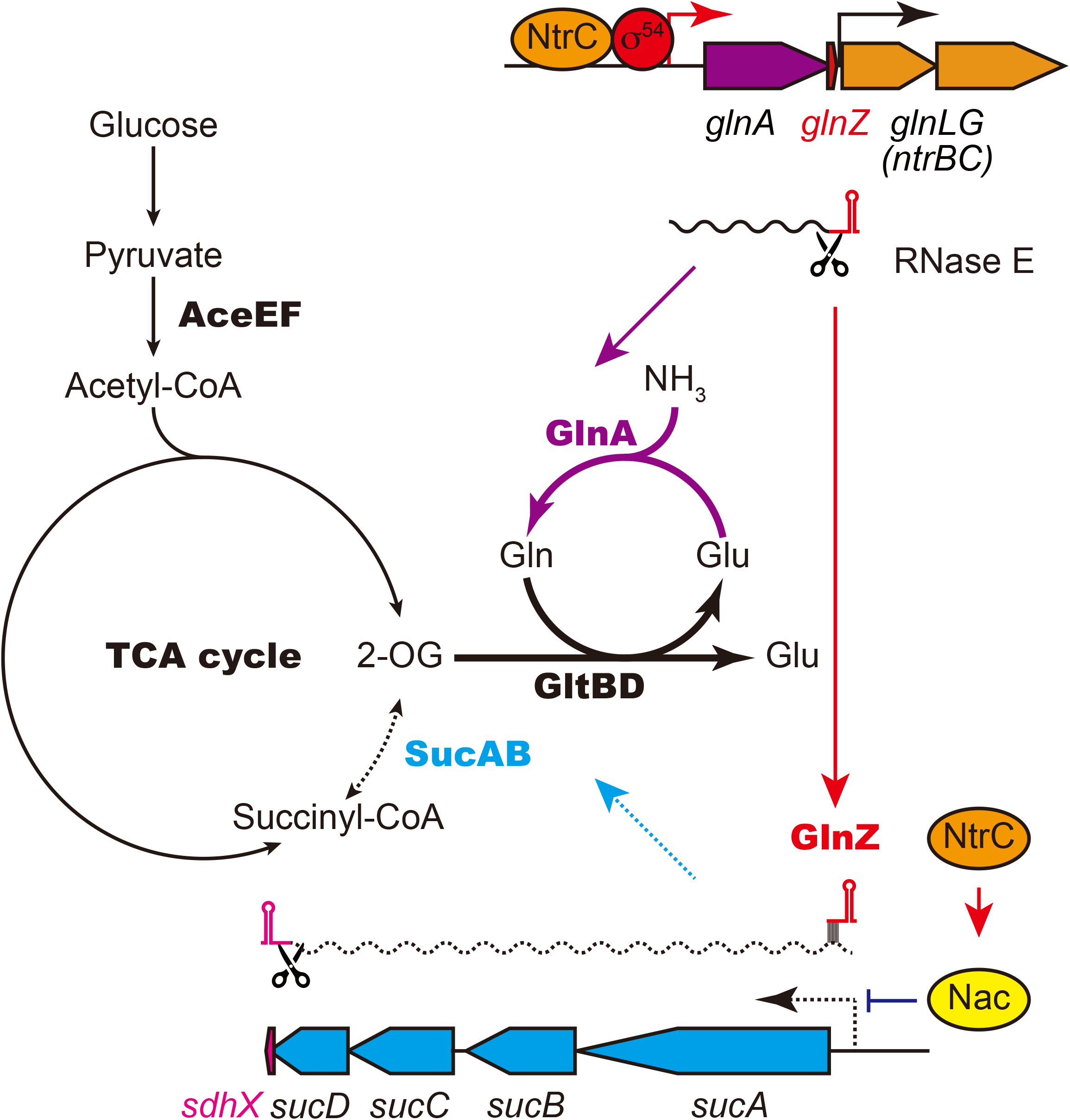
GlnA and GlnZ expressed from *glnA* mRNA facilitate nitrogen assimilation independently. The expression of GlnA and GlnZ is induced by the two-component system NtrBC upon nitrogen limitation. The GS-GOGAT pathway assimilates ammonia using 2-OG as the carbon skeleton. GlnZ sRNA is released from the 3’
sUTR of *glnA* mRNA by RNase E to repress the expression of SucA at the post-transcriptional level. In *E. coli*, the transcriptional regulator Nac, whose expression is activated by NtrC, represses the transcription of *sucABCD* operon in parallel. The repression of OGDH upon nitrogen limitation results in the accumulation of 2-OG and redirects the carbon flow from the TCA cycle to the GS-GOGAT pathway.

### Multilayered regulation of nitrogen assimilation pathway

Under the transcriptional control of the NtrBC two-component system, the *glnA* mRNA is induced upon nitrogen limitation and expresses both GS and GlnZ to facilitate nitrogen assimilation. Although the activity of GS is tightly controlled through post-translational modification (Huergo *et al*, 2013), it seems counterintuitive that the expression of GS is strikingly elevated in the presence of its metabolic product. By contrast, Gram-positive bacteria control the expression of GS through direct interaction between GS and the transcriptional regulator GlnR (Travis *et al*, 2022). Cyanobacteria and archaea have each evolved unique mechanisms to post-transcriptionally regulate the global nitrogen cycle and its metabolism (Prasse & Schmitz, 2018; Moeller *et al*, 2021). The sRNA_154_ of metanogenic archaea *Methanosarcina mazei* controls the expression of GS and P_II_-like signal transduction protein (Prasse *et al*, 2017). In *Synechocystis*, the global nitrogen regulator NtcA upregulates *glnA* in response to increased 2-OG levels and downregulates *gifA* and *gifB* encoding the GS inactivating factors IF7 and IF17, respectively (García-Domínguez *et al*, 2000). NtcA also activates the expression of NsiR4 sRNA, which represses the *gifA* gene at the post-transcriptional level (Klähn *et al*, 2015). The expression of *gifB* is largely enhanced by a Gln-binding riboswitch in its 5’
sUTR, enabling rapid negative feedback of GS activity in response to its product (Klähn *et al*, 2018).

This study reveals that *S. enterica* and *E. coli* utilize the GlnZ sRNA derived from the *glnA* mRNA to control the metabolic branch point at the post-transcriptional level. A Gram-positive bacterium, *Corynebacterium glutamicum*, also tunes the bottleneck in the TCA cycle to produce high levels of Glu by inhibiting the OGDH activity with a 15-kDa protein OdhI depending on its phosphorylation state (Niebisch *et al*, 2006). Surprisingly, in *V. cholerae*, the VcdRP dual-functional sRNA represses the PTS-coding mRNAs while the 29-aa small protein VcdP activates citrate synthase GltA to accumulate 2-OG and Glu (Venkat *et al*, 2021). Altogether, bacteria and archaea employ various RNAs and proteins to regulate the crucial metabolic pathways. Since deletion of *glnZ* and *nac* did not fully restore the SucA level in *E. coli* (Fig. 4B), additional factors may redundantly control the metabolic balance between carbon and nitrogen.

### Conservation and variation of the *glnAZLG* locus and the GlnZ regulon

The *E. coli glnA* 3’
sUTR can be classified into three classes. We demonstrate that the difference in the *glnA* 3’UTRs does not affect the expression of downstream *glnLG* genes (Fig. 2D), and thus the regulon of NtrBC two-component system. The *glnAZLG* genetic organization is conserved in *Enterobacteriaceae*, but *glnA* and *glnLG* (*ntrBC*) genes are often separated in the other γ-Proteobacteria. For instance, *Pseudomonas aeruginosa* harbors four ORFs between *glnA* and *ntrBC*. Moreover, a RpoN-dependent sRNA NrsZ has been found downstream of *ntrBC*, which regulates rhamnolipid biosynthesis and motility (Wenner *et al*, 2014). In the corresponding locus, *Salmonella* carries the *yshB* gene (Hemm *et al*, 2008), which encodes a small protein involved in the intracellular replication of *Salmonella* (Bomjan *et al*, 2019). However, *yshB* contains its own promoter and is not induced in nitrogen limiting conditions.

The GlnZ seed region is highly conserved to regulate *sucA* and *glnP* in common but is located at different positions in the *glnA* 3’UTR in *S. enterica* and *E. coli*. The repression of GlnHPQ ABC transporter by GlnZ may form a negative feedback loop to maintain the intracellular concentration of Gln. *E. coli* GlnZ might regulate other species-specific genes using its unique sequence, as in the case of *E. coli* SdhX repressing *katG* (de Mets *et al*, 2019; Miyakoshi *et al*, 2019). By contrast, *aceE* is repressed by GlnZ in *E. coli*, but *S. enterica* GlnZ1 is deficient in this regulation due to the extra G adjacent to the seed region (Fig. 3D). We further identified the *S. enterica*-specific target *deoD* encoding the purine nucleoside phosphorylase, with which base-pairing interaction involves the extra G nucleotide of GlnZ1 (Supplementary Fig. S1B). Notably, all *E. coli* strains do not have the G nucleotide adjacent to the seed region, but 11 out of 55 *E. fergusonii* strains possess the 41st G nucleotide instead of U in the *glnA* 3’UTRs (Supplementary Figure S2). *E. fergusonii* GlnZ resembles that of *E. coli* O157 with a few nucleotide substitutions, and as expected, the U41G mutation in the O157 GlnZ reduced its activity on *aceE* (Supplementary Figure S2). We speculate that the base-pairing between *pdhR-aceEF* mRNA and GlnZ sRNA is beneficial for *E. coli* but is subject to trade-offs among the other targets.

### Both OGDH and PDH mRNAs are targeted by GlnZ and RNase III in *E. coli*

*sucA* and *aceE* encode components of analogous multienzyme complexes OGDH and PDH, respectively. These enzymes associate with the common E3 subunit, Lpd, and are competitive for the cofactor HS-CoA to produce succinyl-CoA and acetyl-CoA, respectively (Shimada *et al*, 2021). Moreover, these *E. coli* GlnZ target mRNAs contain relatively long stem-loop structures recognized by RNase III in *E. coli* (Cunningham & Guest, 1998; Gordon *et al*, 2017), while GlnZ is also a substrate of RNase III (Walling *et al*, 2022).

Intriguingly, we found that SucA is downregulated and AceE is upregulated in the *rnc14* mutant (Supplementary Figure S3), showing RNase III-mediated regulation in opposite directions. RNase III cleaves the *sucABCD* transcript at three sites upstream of the GlnZ target site (Cunningham & Guest, 1998), but the RNase III cleavage site on the *pdhR-aceEF* mRNA is located downstream of the GlnZ target site (Gordon *et al*, 2017). Thus, the processed *aceEF* mRNA becomes resistant to GlnZ. Nonetheless, our translational fusion constitutively transcribing the *pdhR-aceE* intergenic region was sensitive to GlnZ, indicating that GlnZ is able to base-pair with the *pdhR-aceEF* mRNA (Fig. 4C). Further research will elucidate how RNase III and RNase E mediate the interplay between these important metabolic mRNAs.

### Difference between mRNA-derived sRNAs and dual-function sRNAs

mRNA-derived sRNAs have emerged from diverse regions of mRNAs (Adams & Storz, 2020). In enterobacteria, 3’
sUTR often binds with RNA chaperones Hfq and ProQ, thus serving as a reservoir of functional sRNAs (Miyakoshi *et al*, 2015b; Ponath *et al*, 2022). Surprisingly, regardless of Hfq functionality, 3’UTR-derived sRNAs are also found in Gram-positive bacteria (Fuchs *et al*, 2021; Desgranges *et al*, 2022; Lalaouna *et al*, 2019). Moreover, several 5’UTR-derived and ORF-internal RNAs have been discovered in *S. enterica* and *E. coli* (Adams *et al*, 2021; Matera *et al*, 2022). Wherever sRNAs are derived, their parental mRNAs share identical nucleotide sequences with the sRNAs and thus can function as post-transcriptional regulators. Likewise, dual-function sRNAs are actually mRNAs that encode small peptides and base-pair with target RNAs *in trans* (Raina *et al*, 2018). The two events on dual-function sRNAs are mutually exclusive depending on translation efficiency, RNA stability, and temperature (Aoyama *et al*, 2021; Raina *et al*, 2022; Balasubramanian & Vanderpool, 2013; Aoyama *et al*, 2022). Moreover, post-transcriptional regulation via mRNA-mRNA interactions is not rare in Gram-positive bacteria (Liu *et al*, 2015; Chen *et al*, 2015; Ignatov *et al*, 2020; Mediati *et al*, 2022).

Nonetheless, whether the processing is a prerequisite for the mRNA-derived sRNAs remained obscure. This study first demonstrated that GlnZ sRNA is independent of its parental mRNA but acts in the same biological pathway as the translated protein. This is not simply attributable to sRNA maturation through RNase E cleavage as in the case of ArcZ (Chao *et al*, 2017) since the precursor GlnZ sRNA but not the *glnA* mRNA was capable of target regulation (Fig. 6C). We suggest that the translation of *glnA* mRNA interferes with the base-pairing interaction because the seed region is close to the stop codon in *E. coli* K-12 (Fig. 6A). That said, this study does not exclude the possibility that a parental mRNA itself also interacts with its target mRNA in Gram-negative bacteria. The spacer length between the stop codon and the seed region varies among species and depends on genomic loci. The highly conserved CpxQ sRNA contains two seed regions, R1 and R2, downstream of the *cpxP* stop codon (Chao & Vogel, 2016), the latter of which does not overlap with the RNase E cleavage site. In contrast, the NarS sRNA is processed at a conserved RNase E cleavage site far upstream of the *narK* stop codon, but *E. coli* NarS is much larger than *Salmonella* NarS due to a 170-nt insertion sequence downstream of the *narK* stop codon (Wang *et al*, 2020). It needs to be explored further how the prokaryotic gene expression system coordinates translation, processing, and base-pairing events occurring on a single mRNA molecule.

## MATERIALS AND METHODS

### Bacterial strains

*Salmonella enterica* serovar Typhimurium strain SL1344 (JVS-1574) and *E. coli* strain BW25113 were used as wild-type strains. The strains used in this study are listed in Supplementary Table S2. Bacterial cells were grown at 37°C with reciprocal shaking at 180 rpm in LB Miller medium (BD Biosciences) or MOPS minimal medium (Neidhardt *et al*, 1974). As a carbon source, MOPS minimal medium was supplemented with 0.2% glucose or 20 mM sodium pyruvate. Where appropriate, the nitrogen source of the MOPS minimal medium was substituted by 0.01% of ammonium chloride or 5 mM glutamine (prepared freshly to avoid spontaneous hydrolysis) for 0.1% ammonium chloride. For SL1344 and its derivatives, 40 µM histidine was added into MOPS minimal medium, which does not interfere with GS expression. Optical density (OD_600_) was automatically monitored throughout the growth every 10 min using OD-Monitor C&T (TAITEC). Where appropriate, media were supplemented with antibiotics at the following concentrations: 100 µg/ml ampicillin (Ap), 50 µg/ml kanamycin (Km), and 20 µg/ml chloramphenicol (Cm).

### Plasmid construction

A list of all plasmids and oligonucleotides used in this study can be found in Supplementary Tables S3 and S4. Constitutive expression plasmids, arabinose-inducible pBAD expression plasmids, and translational fusion plasmids were derived from pZE12-luc, pKP8-35, and pXG30-sf, respectively (Urban & Vogel, 2007; Papenfort *et al*, 2006; Corcoran *et al*, 2012). Single-nucleotide mutations were introduced by inverse PCR using KOD-Plus-Neo DNA polymerase (Toyobo) and overlapping primer pairs (Supplementary Table S4) followed by DpnI digestion. The nucleotide sequences of the plasmid inserts can be found in Supplementary Table S5.

### Strain construction

The *glnZ* deletion strains were constructed by the λ Red system (Datsenko & Wanner, 2000). The *glnA* 3’UTR was deleted using pKD4 as a template and primer pairs, MMO-0371/0372 for *Salmonella* and MMO-0401/0402 or MMO-0401/0583 for *E. coli*, respectively. The resulting Km resistant strains were confirmed by PCR, and the mutant loci were transduced into appropriate genetic backgrounds by P22 and P1 phages in *Salmonella* and *E. coli*, respectively. The temperature-sensitive plasmid pCP20 expressing FLP recombinase was used to eliminate the resistance gene from the chromosome.

The *ams-1* locus encoding a temperature-sensitive RNase E mutant and Δ*hfq*::*cat* were transduced from strain TM151 (Morita *et al*, 2006) and TM587 (Morita *et al*, 2005) into appropriate genetic backgrounds by P1 phage, respectively.

The *glnZ* locus in BW25113 was substituted for *glnZ* from O157 or O111 by scar-less mutagenesis using a two-step λRed system (Blank *et al*, 2011). The DNA fragment containing a Cm^R^ resistance marker and a I-SceI recognition site was amplified with primer pairs MMO-0401/0451 using pWRG100 plasmid as a template and was integrated into the BW25113 chromosome by λ Red recombinase expressed from pKD46. The resultant Cm^R^ mutant was transformed by pWRG99, and the heterogeneous *glnZ* allele from O157 or O111 was amplified with MMO-0786/0787 and integrated into the chromosome by λ Red recombinase expressed from pWRG99. The resultant recombinants were selected on LB agar plate supplemented with Ap and 2 µg/ml of anhydrotetracycline to express I-SceI endonuclease. The successful recombinants were confirmed by Cm sensitivity, PCR, and sequencing.

The 3xFLAG epitope tag at the C-terminus of *ntrC* was amplified with primer pairs MMO-0757/0758 using pSUB11 (Uzzau *et al*, 2001) as a template and was introduced into the chromosome by the lambda Red system. The resulting Km resistant strains were confirmed by PCR, and the mutant locus was transduced into appropriate genetic backgrounds by P1 phage.

### GFP fluorescence quantification

*E. coli* Δ*glnZ* transformants harboring a combination of the translational fusions and the GlnZ constitutive expression plasmids (Supplementary Tables S3, S5, and S7) were inoculated from single colonies in 500 µL LB medium containing Ap and Cm in 96-well deep well plates (Thermo Scientific) and were grown overnight at 37°C with rotary shaking at 1,200 rpm in DWMax M-BR-032P plate shaker (Taitec). The 100-µl overnight cultures were dispensed into 96-well optical bottom black microtiter plates (Thermo Scientific). OD_600_ and fluorescence (excitation at 485 nm and emission at 535 nm with dichroic mirror of 510 nm, fixed gain value of 50) were measured using Spark plate reader (Tecan). The relative fluorescence units (RFU) were calculated by subtracting the autofluorescence of bacterial cells without *gfp* grown in the same condition and normalized by OD_600_.

### Northern blot

Total RNA was isolated using the TRIzol reagent (Invitrogen), treated with TURBO DNase (Invitrogen), and precipitated with cold ethanol. RNA was quantified using NanoDrop One (Invitrogen). Total RNA (5 µg) was separated by gel electrophoresis on 6% or 8% polyacrylamide/7 M urea gels in 1×TBE buffer for 2.5 h at 300 V using Biometra Eco-Maxi system (Analytik-Jena). DynaMarker RNA Low II ssRNA fragment (BioDynamics Laboratory) was used as a size marker. RNA was transferred from the gel onto Hybond N+ membrane (Cytiva) by electroblotting for 1 h at 50V using the same device. The membrane was crosslinked with transferred RNA by 120 mJ/cm^2^ UV light, incubated for prehybridization in Rapid-Hyb buffer (Cytiva) at 42°C for 1 h, and then incubated for hybridization with a [^32^P]-labeled probe (Supplementary Table S4) at 42°C overnight. The membrane was washed in three 15-min steps in 5× SSC/0.1% SDS, 1× SSC/0.1% SDS and 0.5× SSC/0.1% SDS buffers at 42°C. Signals were visualized on Typhoon FLA7000 scanner (GE Healthcare) and quantified using Image Quant TL software (GE Healthcare).

For northern blot analysis of mRNA, total RNA samples (2 µg) were resolved by 1.5 % agarose gel electrophoresis in the presence of formaldehyde and blotted onto a Nylon Membrane, positively charged (Roche). Prestein Marker for RNA high (BioDynamics Laboratory) was used as a size marker. The RNAs were visualized by using a detection system with digoxigenin (DIG) (Roche) and then captured using the imaging system ChemiDoc XRS Plus (BioRad). The antisense RNA probes corresponding to the 3’
s-end portion of *glnA* CDS were prepared by the DIG RNA Labeling Kit (Roche).

### Western blot

Bacteria culture was collected by centrifugation for 5 min at 5,000 *g* at 4°C, and the pellet was dissolved in 1x protein loading buffer to a final concentration of 0.01 OD/µl. After heating for 5 min at 95°C, 0.002 OD of whole-cell samples were separated on 7.5% TGX gels (Bio-Rad). Proteins were transferred onto Hybond P PVDF 0.2 membranes (Cytiva), and membranes were blocked for 10 min in Bullet Blocking One (Nacalai Tesque, Kyoto, Japan) and were incubated for 1 h at RT or overnight at 4°C with rabbit polyclonal α-SucA (1:10000, Tanpaku Seisei Co., Gunma, Japan), α-SucB (1:10000, Tanpaku Seisei Co., Gunma, Japan), α-AceE (1:10000, Tanpaku Seisei Co., Gunma, Japan), α-GlnA (1:5000, raised against a synthetic peptide CAHQVNAEFFEEGKMFDGSS, Sigma-Aldrich), α-GroEL (1:10000, Sigma-Aldrich #G6532) and mouse monoclonal α-FLAG (1:5000, Sigma-Aldrich #F1804) diluted in Bullet Blocking One. Membranes were washed three times for 15 min in 1×TBST buffer at RT. Then membranes were incubated for 1 h at RT with secondary anti-mouse or anti-rabbit HRP-linked antibodies (Cell Signaling Technology #7076 or # 7074; 1:5000) diluted in Bullet Blocking One and were washed three times for 15 min in 1×TBST buffer. Chemiluminescent signals were developed using Amersham ECL Prime reagents (Cytiva), visualized on LAS4000 (GE Healthcare), and quantified using Image Quant TL software.

### RNA-seq analysis

Biological duplicates of *S*. Typhimurium SL1344 strain harboring pKP-8-35 or pBAD-GlnZ1 plasmids were grown to exponential phase (OD_600_ ∼1.0) in LB medium. sRNA expression was induced by adding 0.2% L-arabinose for 10 min. Cell cultures were immediately mixed with 20% (v/v) of stop solution (95% ethanol, 5% phenol) and snap-frozen in liquid nitrogen. Total RNA was isolated using the TRIzol reagent (Invitrogen) and treated with TURBO DNase (Invitrogen). Ribosomal RNA was depleted using NEBNext rRNA Depletion Kit (Bacteria), and RNA integrity was confirmed using TapeStation (Agilent). Directional cDNA libraries were prepared using the NEBNext Ultra II Directional RNA Library Prep Kit for Illumina (#7760L). The libraries were sequenced using NovaSeq 6000 platform in the single-read mode for 101 cycles.

Reads were mapped to the *S*. Typhimurium SL1344 reference genome (GenBank accession number FQ312003.1) using bowtie2 version 2.3.2, and HTSeq version 0.5.3p3 was used to generate the count matrix. Reads mapping to annotated coding sequences were counted, normalized (CPM), and transformed (log_2_ Fold Change). Differential expression between the conditions was tested using edgeR version 3.28.0. Genes with a log_2_ fold change >2.0 and FDR adjusted *p*-value < 0.01 were defined as differentially expressed. The RNA-seq data have been deposited in DDBJ DRA under accession number DRA012682.

## ACKNOWLEDGMENTS

The authors would like to thank Jörg Vogel for critical reading, Maxence Lejars, Takeshi Kanda, and Nozomu Obana for discussion and help with preparing figures, Natsuko Shirai for technical assistance, and NBRP-*E*.*coli* at NIG for providing *E. coli* strains. This work was partly performed in the Cooperative Research Project Program of the Medical Institute of Bioregulation, Kyushu University.

