## Supplementary Information for "Glutamine synthetase mRNA releases sRNA from its 3’UTR to regulate carbon/nitrogen metabolic balance"

### This supplement contains:

Supplementary Figures S1 to S3

Supplementary Tables S1 to S7

Figure S1. Post-transcriptional regulation of *Salmonella glnP* and *deoD*

Figure S2. Post-transcriptional regulation of *aceE* by $\mathrm{GlnZ_{O157}}$

Figure S3. Expression of OGDH and PDH components in *E. coli rnc14* mutant

Table S1. Genes downregulated upon GlnZ1 overexpression in *S.* Typhimurium SL1344

Table S2. Bacterial strains used in this study

Table S3. Plasmids used in this study

Table S4. DNA oligonucleotides used in this study

Table S5. Inserts of GlnZ mutant plasmids

Table S6. Details of GFP fusion plasmids

Table S7. Inserts of GFP fusion plasmids

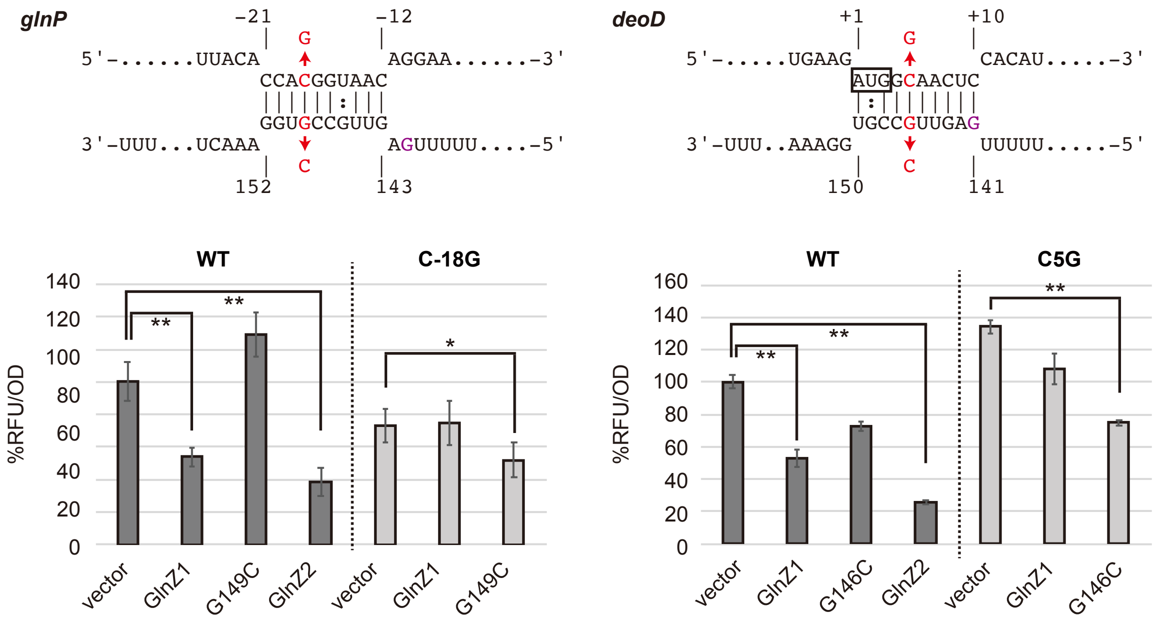


**Fig. S1. Post-transcriptional regulation of *Salmonella glnP* and *deoD*.** Predicted interactions of GlnZ1 with the target mRNAs are shown above the bar graphs. The nucleotide numbers relative to the start codon of the target mRNAs and the stop codon of *glnA* are shown above and below the nucleotide sequences, respectively. The mutated nucleotides were indicated in red letters, and the extra G nucleotide found in *Salmonella* GlnZ1 is shown by purple font. *E. coli* Δ*glnZ* strain was transformed by GFP translational fusion plasmids along with pJV300 control vector or GlnZ expression plasmids (Supplementary Tables S5-7). Mean relative fluorescence units (RFU) normalized by $OD_{600}$ calculated from biological replicates (n>3) are presented with standard deviation in percentage relative to the vector control. Statistical significance was calculated using one-way ANOVA and denoted as follows: ** $p<0.005$, * $p<0.05$.

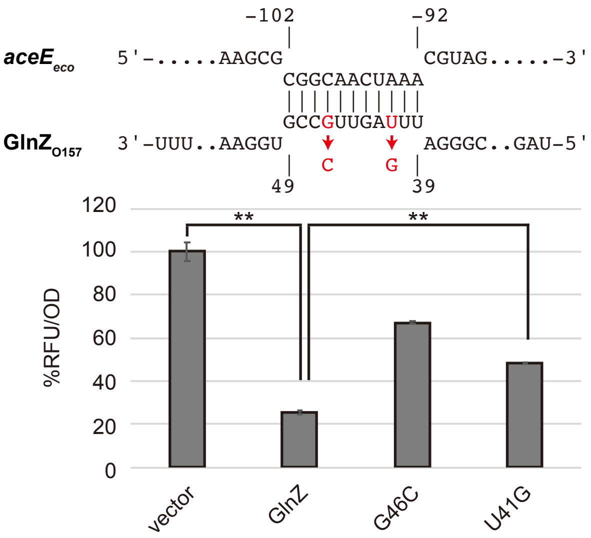


```
CLUSTAL O(1.2.4) multiple sequence alignment                41  seed            terminator

RHB02-C14          tagttgaagttgtactacccggcgctaagaaggccgggatttagttgccgtggaaactttcagcccatctctgcatgggctttttctccatcaattctc-   100
RHB02-C15          tagttgaagttgtactacccggcgctaagaaggccgggatttagttgccgtggaaactttcagcccatctctgcatgggctttttctccatcaattctc-   100
RHB02-C18          tagttgaagttgtactacccggcgctaagaaggccgggatttagttgccgtggaaactttcagcccatctctgcatgggctttttctccatcaattctc-   100
RHB02-C22          tagttgaagttgtactacccggcgctaagaaggccgggatttagttgccgtggaaactttcagcccatctctgcatgggctttttctccatcaattctc-   100
RHB28-C13          tagttgaagttgtactacccggcgctaagaaggccgggatttagttgccgtggaaactttcagcccatctctgcatgggctttttctccatcaattctc-   100
RHB30-C07          tagttgaagttgtactacccggcgctaagaaggccgggatttagttgccgtggaaactttcagcccatctctgcatgggctttttctccatcaattctc-   100
RHB38-C01          tagttgaagttgtactacccggcgctaagaaggccgggatttagttgccgtggaaactttcagcccatctctgcatgggctttttctccatcaattctc-   100
5zf15-2-1          tagttgaagttgtactacccggtgctaagaaggccgggatttagttgccgtggaaactttcagcccatctctgcatgggctttttctccgtaacttctt-   100
6S41-1             tagttgaagttgtactacccggtgctaagaaggccgggatttagttgccgtggaaactttcagcccatctctgcatgggctttttctccgtaacttctt-   100
40A                tagttgaagttgtactacccggtgctaagaaggccgggatttagttgccgtggaaactttcagcccatctctgcatgggctttttctccgtaacttctt-   100
ATCC35470          tagttgaagttgtactacccggtgctaagaaggccgggatttagttgccgtggaaactttcagcccatctctgcatgggctttttctccgtaacttctt-   100
ATCC35471          tagttgaagttgtactacccggtgctaagaaggccgggatttagttgccgtggaaactttcagcccatctctgcatgggctttttctccgtaacttctt-   100
ATCC35473          tagttgaagttgtactacccggtgctaagaaggccgggatttagttgccgtggaaactttcagcccatctctgcatgggctttttctccgtaacttctt-   100
EF31               tagttgaagttgtactacccggtgctaagaaggccgggatttagttgccgtggaaactttcagcccatctctgcatgggctttttctccgtaacttctt-   100
FDAARGOS_1438      tagttgaagttgtactacccggtgctaagaaggccgggatttagttgccgtggaaactttcagcccatctctgcatgggctttttctccgtaacttctt-   100
RHB03-C23          tagttgaagttgtactacccggtgctaagaaggccgggatttagttgccgtggaaactttcagcccatctctgcatgggctttttctccgtaacttctt-   100
RHB10-C04          tagttgaagttgtactacccggtgctaagaaggccgggatttagttgccgtggaaactttcagcccatctctgcatgggctttttctccgtaacttctt-   100
RHB10-C23          tagttgaagttgtactacccggtgctaagaaggccgggatttagttgccgtggaaactttcagcccatctctgcatgggctttttctccgtaacttctt-   100
RHB17-C11          tagttgaagttgtactacccggtgctaagaaggccgggatttagttgccgtggaaactttcagcccatctctgcatgggctttttctccgtaacttctt-   100
RHB25-C10          tagttgaagttgtactacccggtgctaagaaggccgggatttagttgccgtggaaactttcagcccatctctgcatgggctttttctccgtaacttctt-   100
RHB32-C05          tagttgaagttgtactacccggtgctaagaaggccgggatttagttgccgtggaaactttcagcccatctctgcatgggctttttctccgtaacttctt-   100
RHB32-C09          tagttgaagttgtactacccggtgctaagaaggccgggatttagttgccgtggaaactttcagcccatctctgcatgggctttttctccgtaacttctt-   100
RHB32-C16          tagttgaagttgtactacccggtgctaagaaggccgggatttagttgccgtggaaactttcagcccatctctgcatgggctttttctccgtaacttctt-   100
RHB32-C22          tagttgaagttgtactacccggtgctaagaaggccgggatttagttgccgtggaaactttcagcccatctctgcatgggctttttctccgtaacttctt-   100
RHB33-C04          tagttgaagttgtactacccggtgctaagaaggccgggatttagttgccgtggaaactttcagcccatctctgcatgggctttttctccgtaacttctt-   100
RHB33-C07          tagttgaagttgtactacccggtgctaagaaggccgggatttagttgccgtggaaactttcagcccatctctgcatgggctttttctccgtaacttctt-   100
RHB38-C04          tagttgaagttgtactacccggtgctaagaaggccgggatttagttgccgtggaaactttcagcccatctctgcatgggctttttctccgtaacttctt-   100
RHB42-C01          tagttgaagttgtactacccggtgctaagaaggccgggatttagttgccgtggaaactttcagcccatctctgcatgggctttttctccgtaacttctt-   100
RHB42-C13          tagttgaagttgtactacccggtgctaagaaggccgggatttagttgccgtggaaactttcagcccatctctgcatgggctttttctccgtaacttctt-   100
RHB42-C24          tagttgaagttgtactacccggtgctaagaaggccgggatttagttgccgtggaaactttcagcccatctctgcatgggctttttctccgtaacttctt-   100
RHB24-C06          tagttgaagttgtactacccggtgctaagaaggccgggatttagttgccgtggaaactttcagcccatctctgcatgggctttttctccgtcaattctt-   100
RHB24-C08          tagttgaagttgtactacccggtgctaagaaggccgggatttagttgccgtggaaactttcagcccatctctgcatgggctttttctccgtcaattctt-   100
ATCC35469          tagttgaagttgtactacccggcgc-aacaacgccgggatttagttgccgtggaaactttcagcccatctctgcatgggctttttctccgtcaattctct   100
EF44               tagttgaagttgtactacccggcgc-aacaacgccgggatttagttgccgtggaaactttcagcccatctctgcatgggctttttctccgtcaattctct   100
EF91               tagttgaagttgtactacccggcgc-aacaacgccgggatttagttgccgtggaaactttcagcccatctctgcatgggctttttctccgtcaattctct   100
EFCF056            tagttgaagttgtactacccggcgc-aacaacgccgggatttagttgccgtggaaactttcagcccatctctgcatgggctttttctccgtcaattctct   100
FDAARGOS_1032      tagttgaagttgtactacccggcgc-aacaacgccgggatttagttgccgtggaaactttcagcccatctctgcatgggctttttctccgtcaattctct   100
FDAARGOS_1499      tagttgaagttgtactacccggcgc-aacaacgccgggatttagttgccgtggaaactttcagcccatctctgcatgggctttttctccgtcaattctct   100
RHB19-C05          tagttgaagttgtactacccggcgc-aacaacgccgggatttagttgccgtggaaactttcagcccatctctgcatgggctttttctccgtcaattctct   100
RHB31-C13          tagttgaagttgtactacccggcgc-aacaacgccgggatttagttgccgtggaaactttcagcccatctctgcatgggctttttctccgtcaattctct   100
RHB38-C07          tagttgaagttgtactacccggcgc-aacaacgccgggatttagttgccgtggaaactttcagcccatctctgcatgggctttttctccgtcaattctct   100
RHB41-C15          tagttgaagttgtactacccggcgc-aacaacgccgggatttagttgccgtggaaactttcagcccatctctgcatgggctttttctccgtcaattctct   100
RHB41-C20          tagttgaagttgtactacccggcgc-aacaacgccgggatttagttgccgtggaaactttcagcccatctctgcatgggctttttctccgtcaattctct   100
O157               tagttgaagttgtactacccggcgc-aacaacgccgggatttagttgccgtggaaactttcagcccatctctgcatgggctttttctccgtcaattctct   100
HNCF11W            tagttgaagttgtactacccggcgc-aacaacgccgggatttagttgccgtggaaactttcagcccatctctgcatgggctttttctccatcaattctct   100
EF20JDJ4045        tagttgaagttgtactacccggcgc-aacaacgccgggattgagttgccgtggaaactttcagcccatctctgcatgggctttttctccataacttcttt   100
RHB18-C03          tagttgaagttgtactacccggcgc-aacaacgccgggattgagttgccgtggaaactttcagcccatctctgcatgggctttttctccataacttcttt   100
RHB18-C04          tagttgaagttgtactacccggcgc-aacaacgccgggattgagttgccgtggaaactttcagcccatctctgcatgggctttttctccataacttcttt   100
RHB18-C08          tagttgaagttgtactacccggcgc-aacaacgccgggattgagttgccgtggaaactttcagcccatctctgcatgggctttttctccataacttcttt   100
RHB26-C03          tagttgaagttgtactacccggcgc-aacaacgccgggattgagttgccgtggaaactttcagcccatctctgcatgggctttttctccataacttcttt   100
RHB41-C23          tagttgaagttgtactacccggcgc-aacaacgccgggattgagttgccgtggaaactttcagcccatctctgcatgggctttttctccataacttcttt   100
RHB23-C01          tagttgaagttgtactacccggcgc-aacaacgccgggattgagttgccgtggaaactttcagcccatctctgcatgggctttttctccgtaacttcttt   100
RHB23-C02          tagttgaagttgtactacccggcgc-aacaacgccgggattgagttgccgtggaaactttcagcccatctctgcatgggctttttctccgtaacttcttt   100
RHB26-C09          tagttgaagttgtactacccggcgc-aacaacgccgggattgagttgccgtggaaactttcagcccatctctgcatgggctttttctccgtaacttcttt   100
RHB28-C21          tagttgaagttgtactacccggcgc-aacaacgccgggattgagttgccgtggaaactttcagcccatctctgcatgggctttttctccgtaacttcttt   100
RHB31-C17          tagttgaagttgtactacccggcgc-aacaacgccgggattgagttgccgtggaaactttcagcccatctctgcatgggctttttctccgtaacttcttt   100
                   ********************** ** ** ** ********* *********************************************** * * ****
```

**Fig. S2. Post-transcriptional regulation of *aceE* by GlnZ$_{O157}$.** (A) *E. coli* Δ*glnZ* strain was transformed by pXG30sf-*aceE*$_{eco}$ along with pJV300 control vector or GlnZ$_{O157}$ expression plasmids (Supplementary Tables S5-7). Mean relative fluorescence units (RFU) normalized by OD$_{600}$ calculated from biological replicates (n>3) are presented with standard deviation in percentage relative to the vector control. Statistical significance was calculated using one-way ANOVA and denoted as follows: ** $p$<0.005. (B) Alignment of *E. fergusonii* GlnZ from 55 strains along with GlnZ$_{O157}$. GlnZ$_{O157}$ is shown in bold font. Nucleotides of *E. fergusonii* GlnZ different from those of GlnZ$_{O157}$ are highlighted in yellow and indicated in red font.

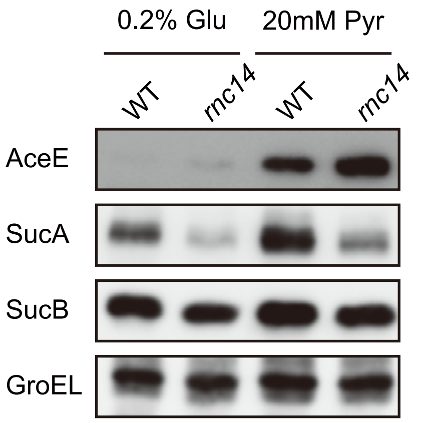


**Fig. S3. Expression of OGDH and PDH components in *E. coli rnc14* mutant.** *E. coli* WT and *rnc14* strains were grown to exponential phase (OD600 ~0.5) in MOPS minimal medium containing 0.2% glucose or 20 mM pyruvate as the carbon source and 0.1% ammonium as the nitrogen source. The expression of AceE, SucA and SucB, and GroEL was analyzed by western blot.

**Table S1. Genes downregulated upon GlnZ1 overexpression in *S*. Typhimurium SL1344.**

| **Name** | **$\log_2$ fold change** | ***p* value** | **FDR** |
|---|---|---|---|
| SraB | 3.02 | 3.4E-05 | 3.2E-03 |
| SL1344_4248 | 2.84 | 6.4E-11 | 1.5E-07 |
| *asnA* | 2.71 | 3.6E-09 | 2.8E-06 |
| SL1344_0301 | 2.59 | 1.1E-07 | 4.6E-05 |
| *fabA* | 2.33 | 1.6E-08 | 9.4E-06 |
| *sirC* | 2.31 | 4.6E-06 | 9.1E-04 |
| ***glnP*** | **2.25** | **1.1E-04** | **6.7E-03** |
| ***deoD*** | **2.23** | **1.9E-10** | **3.1E-07** |
| ***glnQ*** | **2.14** | **1.3E-08** | **9.1E-06** |
| *rpsG* | 2.09 | 6.1E-05 | 4.4E-03 |
| *aroQ* | 2.06 | 2.8E-06 | 7.1E-04 |
| *rpsL* | 2.00 | 5.5E-05 | 4.4E-03 |

### Table S2. Bacterial strains used in this study.

| Strain | Relevant markers/ genotype | Reference/ source |
|---|---|---|
| ***S. enterica* subsp. Typhimurium** | | |
| SL1344 | LT2 *hisG* | Laboratory stock |
| Δ*glnZ*::*kan* | SL1344 Δ*glnA* 3´UTR::*kan* | This study |
| Δ*glnZ* | SL1344 Δ*glnA* 3´UTR::FRT | This study |
| ***E. coli*** | | |
| BW25113 | $F^-$ $\lambda^-$ *rrnB3* Δ*lacZ*4787 *hsdR*514 Δ(*araBAD*)567 Δ(*rhaBAD*)568 *rph*-1 | NBPR strain |
| O157 | | Laboratory stock |
| O111 | | Laboratory stock |
| Δ*glnZ*::*kan* | BW25113 Δ*glnZ*::*kan* | This study |
| Δ*glnZ*::*cat* I-SceI | BW25113 Δ*glnZ*:: *cat* I-SceI | This study |
| Δ*glnZT*::*kan* | BW25113 Δ*glnZT*::*kan* | This study |
| Δ*glnZ* | BW25113 Δ*glnZ*::FRT | This study |
| Δ*glnZT* | BW25113 Δ*glnZT*::FRT | This study |
| $glnZ_{O157}$ | BW25113 $glnZ_{O157}$ | This study |
| $glnZ_{O111}$ | BW25113 $glnZ_{O111}$ | This study |
| *ntrC*::3xFLAG | BW25113 *ntrC*::3xFLAG *kan* | This study |
| Δ*glnZ ntrC*:::3xFLAG | BW25113 Δ*glnZ::* FRT *ntrC*::3xFLAG *kan* | This study |
| ΔΔ*glnZ ntrC*:::3xFLAG | BW25113 ΔΔ*glnZ*::FRT *ntrC*::3xFLAG *kan* | This study |
| $glnZ_{O157}$ *ntrC*:::3xFLAG | BW25113 $glnZ_{O157}$ *ntrC*::3xFLAG *kan* | This study |
| $glnZ_{O111}$ *ntrC*:::3xFLAG | BW25113 $glnZ_{O111}$ *ntrC*::3xFLAG *kan* | This study |
| Δ*nac* | BW25113 Δ*nac::kan* (JW1967) | NBPR strain |
| Δ*glnZ* Δ*nac* | BW25113 Δ*glnZ::*FRT Δ*nac::kan* | This study |
| TM151 | W3110 *mlc ams-1*::Tn*10* | (Morita *et al.*, 2006) |
| TM587 | W3110 *mlc* Δ*hfq*::*cat* | (Morita *et al.*, 2005) |
| Δ*glnZ ams-1* | BW25113 Δ*glnZ::*FRT *ams-1*::Tn*10* | This study |
| Δ*glnZ* Δ*hfq* | BW25113 Δ*glnZ::*FRT Δ*hfq*::*cat* | This study |

### Table S3. Plasmids used in this study.

| **Name** | **Relevant fragment** | **Comment** | **Origin / marker** | **Reference** |
|---|---|---|---|---|
| pJV300 | control plasmid | Constitutive expression plasmid | ColE1/ Amp$^{R}$ | (Urban and Vogel, 2007) |
| pP$_{L}$-GlnZ1 | P$_{LlacO}$-*glnZ1* | *Salmonella glnZ1* expression plasmid | ColE1/ Amp$^{R}$ | this study |
| pP$_{L}$-GlnZ1 G149C | P$_{LlacO}$ -*glnZ1* G149C | *Salmonella glnZ1* G149C mutant | ColE1/ Amp$^{R}$ | this study |
| pP$_{L}$-GlnZ1 G146C | P$_{LlacO}$-*glnZ1* G146C | *Salmonella glnZ1* G146C mutant | ColE1/ Amp$^{R}$ | this study |
| pP$_{L}$-GlnZ1 G141U | P$_{LlacO}$-*glnZ1* G141U | *Salmonella glnZ1* G141U mutant | ColE1/ Amp$^{R}$ | this study |
| pP$_{L}$-GlnZ1 G141U/G146C | P$_{LlacO}$-*glnZ1* G141U/G146C | *Salmonella glnZ1* G141U/G146C mutant | ColE1/ Amp$^{R}$ | this study |
| pP$_{L}$-GlnZ2 | P$_{LlacO}$-*glnZ2* | *Salmonella glnZ2* expression plasmid | ColE1/ Amp$^{R}$ | this study |
| pP$_{L}$-GlnZ$_{K12}$ | P$_{LlacO}$- *glnZ* $_{K12}$ | *E. coli glnZ* $_{K12}$ expression plasmid | ColE1/ Amp$^{R}$ | this study |
| pP$_{L}$-GlnZ$_{K12}$ G15C | P$_{LlacO}$- *glnZ* $_{K12}$ G15C | *E. coli glnZ* $_{K12}$ G15C mutant | ColE1/ Amp$^{R}$ | this study |
| pP$_{L}$-GlnZ$_{K12}$ G12C | P$_{LlacO}$- *glnZ* $_{K12}$ G12C | *E. coli glnZ* $_{K12}$ G12C mutant | ColE1/ Amp$^{R}$ | this study |
| pP$_{L}$-GlnZ$_{O157}$ | P$_{LlacO}$-*glnZ* $_{O157}$ | *E. coli glnZ* $_{O157}$ expression plasmid | ColE1/ Amp$^{R}$ | this study |
| pP$_{L}$-GlnZ$_{O157}$ G49C | P$_{LlacO}$-*glnZ* $_{O157}$ G49C | *E. coli glnZ* $_{O157}$ G49C mutant | ColE1/ Amp$^{R}$ | this study |
| pP$_{L}$-GlnZ$_{O157}$ G46C | P$_{LlacO}$- *glnZ* $_{O157}$ G46C | *E. coli glnZ* $_{O157}$ G46C mutant | ColE1/ Amp$^{R}$ | this study |
| pP$_{L}$-GlnZ$_{O157}$ U41G | P$_{LlacO}$- *glnZ* $_{O157}$ U41G | *E. coli glnZ* $_{O157}$ U41G mutant | ColE1/ Amp$^{R}$ | this study |
| pP$_{L}$-GlnZ$_{O111}$ | P$_{LlacO}$-*glnZ* $_{O111}$ | *E. coli glnZ* $_{O111}$ expression plasmid | ColE1/ Amp$^{R}$ | this study |
| pKP8-35 | control plasmid | Arabinose-inducible pBAD plasmid | pBR322/ Amp$^{R}$ | (Papenfort *et al.*, 2006) |
| pBAD-GlnZ1 | P$_{araBAD}$-*glnZ1* | *Salmonella glnZ1* expression plasmid | pBR322/ Amp$^{R}$ | this study |
| pBAD-*glnA sal* | P$_{araBAD}$-*glnA sal* | *Salmonella glnA* expression plasmid | pBR322/ Amp$^{R}$ | this study |
| pBAD-GlnZ+90 *sal* | P$_{araBAD}$- preGlnZ *sal* | *Salmonella* premature *glnZ* expression plasmid | pBR322/ Amp$^{R}$ | this study |
| pBAD-*glnA eco* | P$_{araBAD}$-*glnA eco* | *E. coli glnA* $_{K12}$ expression plasmid | pBR322/ Amp$^{R}$ | this study |
| pBAD-*glnA* mut1 | P$_{araBAD}$-*glnA eco* mut1 | *E. coli glnA* $_{K12}$ mutant in site 1 | pBR322/ Amp$^{R}$ | this study |
| pBAD-*glnA* mut2 | P$_{araBAD}$-*glnA eco* mut2 | *E. coli glnA* $_{K12}$ mutant in site 2 | pBR322/ Amp$^{R}$ | this study |
| pBAD-*glnA* mut3 | P$_{araBAD}$-*glnA eco* mut3 | *E. coli glnA* $_{K12}$ mutant in site 1 and 2 | pBR322/ Amp$^{R}$ | this study |
| pBAD-GlnZ+90 *eco* | P$_{araBAD}$- preGlnZ *eco* | *E. coli* premature *glnZ* (90-nt extension) expression plasmid | pBR322/ Amp$^{R}$ | this study |
| pBAD-GlnZ+30 *eco* | P$_{araBAD}$- GlnZ$_{227}$ *eco* | *E. coli* premature *glnZ* (30-nt extension) expression plasmid | pBR322/ Amp$^{R}$ | this study |
| pBAD-GlnZ+30 *eco* mut1 | P$_{araBAD}$- GlnZ$_{227}$ *eco* mut1 | *E. coli* premature *glnZ* (30-nt extension) mutant in site 1 | pBR322/ Amp$^{R}$ | this study |
| pXG-30sf | P$_{LtetO}$-FLAG::*glmU-glmS::gfp* | Translational sfGFP fusion plasmid for dicistronic targets | pSC101* / Cm$^{R}$ | (Corcoran *et al.*, 2012) |
| pXG-30sf-*sucA sal* | P$_{LtetO}$-FLAG::*sdhB-sucA::gfp* | *S. enterica sdhB-sucA* translational fusion plasmid | pSC101*/ Cm$^{R}$ | this study |
| pXG-30sf-*sucA eco* | P$_{LtetO}$-FLAG::*sdhB-sucA::gfp* | *E. coli sdhB-sucA* translational fusion plasmid | pSC101*/ Cm$^{R}$ | this study |
| pXG-30sf-*deoD sal* | P$_{LtetO}$-FLAG::*deoB-deoD::gfp* | *S. enterica deoBD* translational GFP fusion plasmid | pSC101*/ Cm$^{R}$ | this study |
| pXG-30sf-*glnP sal* | P$_{LtetO}$-FLAG::*glnH-glnP::gfp* | *S. enterica glnHP* translational GFP fusion plasmid | pSC101*/ Cm$^{R}$ | this study |
| pXG-30sf-*aceE sal* | P$_{LtetO}$-FLAG::*pdhR-aceE::gfp* | *S. enterica pdhR-aceE* translational fusion plasmid | pSC101*/ Cm$^{R}$ | this study |
| pXG-30sf-*aceE eco* | P$_{LtetO}$-FLAG::*pdhR-aceE::gfp* | *E. coli pdhR-aceE* translational fusion plasmid | pSC101*/ Cm$^{R}$ | this study |

### Table S4. DNA oligonucleotides used in this study.

| Name | Sequence (5´ -> 3´ direction) | Used for |
|---|---|---|
| **Northern blot** | | |
| MMO-0416 | AAAGTTTCCACGGCAACT | Probe for GlnZ |
| MMO-0417 | ATCCTGGGATGGGCTGAAAG | Probe for GlnZ2 |
| MMO-0418 | AACTCCTGACGCCTTTCACG | Probe for GlnZ1 |
| MMO-0419 | ATGCAGAGATGGGCTACAGA | Probe for GlnZ eco |
| MMO-1056 | ACTACCATCGGCGCTACGGC | Probe for 5S rRNA |
| JVO-2907 | GAAGATTGTTGCCCGGCGATTTG | Probe for SroC sal |
| JVO-5622 | GAAGATTGTTACCCAGCGTATTG | Probe for SroC eco |
| SP6-AS *glnA*sal | ATTGTCGTTAGAACGCGGCTACAATTAATACATAACCTTATGTATCATACAC ATACGATTTAGGTGACACTATAGACGCACGCGGTCATCTTCTTCGCGACGCA GCGCAATATACGCATCGATCGCTTCATCAGTGAACACGCCGCCTGCTTTCAG GAACTCGCGGTCCAGGTCCAGCGCGTTCAGCGCTTCTTCCAGAGAACCCGCT ACCTGTGGGATCTCTTTCGCTTCTTCCGGCGGCAGGTCATACAGGTTTTTGT CCATGGCTTCGCCCG | Probe for *glnA* sal |
| SP6-AS *glnA*eco | ATTGTCGTTAGAACGCGGCTACAATTAATACATAACCTTATGTATCATACAC ATACGATTTAGGTGACACTATAGACGCACGCGGTCATCTTCTTCGCGACGCA GAGCGATGTACGCATCAATTGCTTCGTCAGTGAACACGCCACCGGCTTTCAG GAACTCGCGGTCCAGATCCAGTTCGTTCAGTGCTTCTTCCAGAGAGCCTGCA ACCTGTGGGATCTCTTTCGCTTCTTCTGGCGGCAGGTCATACAGGTTTTTGT CCATGGCTTCGCCCG | Probe for *glnA* eco |
| ***glnA* and GlnZ cloning** | | |
| MMO-0354 | GTTTTTTCTAGATTGTTGGTGGAGAAAAAAG | *glnA*/GlnZ sal |
| MMO-0355 | ATCGTATATTAAAAATCCGACAAATTTC | GlnZ1 sal |
| MMO-0356 | AGTTTTTGAGTTGCCGTGGA | GlnZ2 sal |
| MMO-0384 | ACGGCGACACGGCCAG | *glnA* sal |
| MMO-0386 | GTAGAGTTTGAGCTGTACTACAGC | GlnZ+30 |
| MMO-0399 | ACGGCGACACGGCCAAAATAATTG | *glnA* eco |
| MMO-0405 | AGTGTTTTAGTTGCCGTGG | GlnZ K12 |
| MMO-0406 | GTTTTTTCTAGAATTGACGGAGAAAAAAG | *glnA* /GlnZ eco |
| MMO-0786 | GATGCGTACATCGCTCTG | GlnZ+90 |
| MMO-0787 | CATAAAGCAGTCTCCTGAACA | *glnL* eco from ATG |
| MMO-0788 | ATAGTTGAAGTTGTACTACCC | GlnZ O157/O111 |
| ***glnA* and GlnZ mutagenesis** | | |
| MMO-0361 | GTTGCCCTGGAAACTTTCAGCCCAT | GlnZ sal G149C |
| MMO-0362 | TTTCCAGGGCAACTCAAAAACTCCTG | GlnZ sal G149C |
| MMO-0694 | TGAGTTCCGTGGAAACTTTCAGCC | GlnZ sal G146C |
| MMO-0695 | CCACGGGAACTCAAAAACTCCTGACGC | GlnZ sal G146C |
| MMO-0857 | GTTTTTAGTTGCCGTGGAAACTTTC | GlnZ sal G141U |
| MMO-0858 | GCAACTAAAAACTCCTGACGCCTTTC | GlnZ sal G141U |
| MMO-0859 | GTTTTTAGTTCCCGTGGAAACTTTC | GlnZ sal G141U/G146C |
| MMO-0860 | GGAACTTAAAACTCCTGACGCCTTTC | GlnZ sal G141U/G146C |
| MMO-0411 | GTTGCCCTGGAAACTTTTCGCCTGT | GlnZ K12 G15C |
| MMO-0412 | TTTCCAGGGCAACTAAAACACTGTGCTC | GlnZ K12 G15C |
| MMO-0718 | TTAGTTCCCGTGGAAACTTTTCGCC | GlnZ K12 G12C |
| MMO-0719 | CCACGGGAACTAAAACACTGTGCTCAGT | GlnZ K12 G12C |
| MMO-1168 | GGGATTGAGTTGCCGTGGAAACTTTC | GlnZ O157 U41G |
| MMO-1169 | GCAACTCAATCCCGGCGTTGTTGC | GlnZ O157 U41G |
| MMO-1180 | GTTGCCCTGGAAACTTTCAGCCCATC | GlnZ O157 G49C |
| MMO-1181 | TTTCCAGGGCAACTAAATCCCGGC | GlnZ O157 G49C |
| MMO-1182 | TTAGTTCCCGTGGAAACTTTCAGCC | GlnZ O157 G46C |
| MMO-1183 | CCACGGGAACTAAATCCCGGCGTTG | GlnZ O157 G46C |
| MMO-0617 | GTAGAGCGGGAGCTGTACTACAGCGTCTA | *glnA* K12 RNase E mut2 |
| MMO-0618 | CAGCTCCCGCTCTACCGGATGCGGAG | *glnA* K12 RNase E mut2 |
| MMO-0621 | AAGTGTGGTAGTTGCCGTGGAAACTTTTC | *glnA* K12 RNase E mut1 |
| MMO-0622 | CAACTACCACACTTAGACGCTGTAGTACAG | *glnA* K12 RNase E mut1 |
| **GlnZ target cloning** | | |
| MMO-0325 | GTTTTTATGCATGCATTCAGCGTATTCCG | *sdhB-sucA* GFP fusion |
| MMO-0326 | GTTTTTGCTAGCCAACCAGGCTTTCAAAGC | *sdhB-sucA* GFP fusion |
| MMO-0529 | GTTTTTATGCATCCGATGGACTACGGTAAAA | *deoBDsal* GFP fusion |
| MMO-0530 | GTTTTTGCTAGCCGGCATCAATACGACGT | *deoBDsal* GFP fusion |
| MMO-0594 | GTTTTTATGCATAACGAAATCTACAAAAAATGGT | *glnHP* GFP fusion |
| MMO-0595 | GTTTTTGCTAGCCCAGTCAAACTGCATATGT | *glnHP* GFP fusion |
| MMO-0701 | GTTTTTATGCATGAGATGCTGCCCGCTGGTG | *pdhR-aceE* GFP fusion |
| MMO-0702 | GTTTTTGCTAGCAACACCTTCTTCACGGATGACC | *pdhR-aceE* GFP fusion |
| **GlnZ target mutagenesis** | | |
| MMO-0363 | TATCCAGGGCGAAATACTCGTCATAG | *sucAsal* C-166G |
| MMO-0364 | TTCGCCCTGGATACTAACCACGCATAC | *sucAsal* C-166G |

| | | |
|---|---|---|
| MMO-0369 | TATCCAGGGCGAAGTAAGCATAAAAAAG | *sucAeco* C-38G |
| MMO-0370 | TTCGCCCTGGATACTACCACGCACAG | *sucAeco* C-38G |
| MMO-0691 | ACACCAGGGTAACAGGAACGACATATG | *glnPsal* C-18G |
| MMO-0692 | GTTACCCTGGTGTAAATAGTCAAATGCT | *glnPsal* C-18G |
| MMO-0697 | AGATGGGAACTCCACATATTAATGCAGAAATG | *deoDsal* C5G |
| MMO-0698 | GGAGTTCCCATCTTCAGTTCCCTTTAAATTTG | *deoDsal* C5G |
| MMO-0720 | GCGCGGGAACTAAACGCAGAACCTGTCTTATTAAGC | *aceEsal* C-99G |
| MMO-0721 | TTAGTTCCCGCGCACATTTTTGCGC | *aceEsal* C-99G |
| MMO-0722 | GCGCGGGAACTAAACGTAGAACCTGTCTTATTG | *aceEeco* C-99G |
| MMO-0723 | TTAGTTCCCGCGCTTTTATATGCGC | *aceEeco* C-99G |
| **Lambda Red recombination** | | |
| MMO-0371 | CCCGCACCCGGTAGAGTTTGAGCTGTACTACAGCGTTTAAGTGTAGGCTGGAGCTGCTTC | Sense oligo to construct *Salmonella* Δ*glnZ*::*kan* |
| MMO-0372 | AGATTGTTGGTGGAGAAAAAAGCCCATCCTGGGATGGGCTGGTCCATATGAATATCCTCCTTAG | Antisense oligo to construct *Salmonella* Δ*glnZ*::*kan* |
| MMO-0401 | TCCGCATCCGGTAGAGTTTGAGCTGTACTACAGCGTCTAAGTGTAGGCTGGAGCTGCTTC | Sense oligo to construct *E. coli* Δ*glnZ*::*kan* |
| MMO-0402 | AGAGAATTGACGGAGAAAAAAGCCCATGCAGAGATGGGCTGGTCCATATGAATATCCTCCTTAG | Antisense oligo to construct *E. coli* Δ*glnZ*::*kan* |
| MMO-0451 | AGAGAATTGACGGAGAAAAAAGCCCATGCAGAGATGGGCTCTAGACTATATTACCCTGTT | Antisense oligo to construct *E. coli* Δ*glnZ*::*cat* ISce-I |
| MMO-0583 | TTACGGATAAAAAGCGCGAAGCATCAGAGAATTGACGGAGGGTCCATATGAATATCCTCCTTAG | Antisense oligo to construct *E. coli* Δ*glnZT* ::*kan* |
| MMO-0757 | CAACACCCTGACGCGTAAGTTAAAAGAGCTGGGGATGGAGGACTACAAAGACCATGACGG | *ntrC*::3xFLAG insertion with pSUB13 |
| MMO-0758 | CAATTTGCGCTCAATAATCAATCTTTACACACAAGCTGTGTCCATATGAATATCCTCCTTAG | *ntrC*::3xFLAG insertion with pSUB13 |

**Table S5. Inserts of GlnZ mutant plasmids.**

The modified nucleotides are highlighted in magenta. XbaI sites introduced for cloning were highlighted in yellow.

| **Plasmid** | **Insert from +1 to end of *gcvB* terminator** | **Positions deleted or mutated** |
|---|---|---|
| pP$_{L}$-GlnZ1 | aTCGTATATTAAAAATCCGACAAATTTCGCGTTGCTGCAAGGCAGCAACTGAG<br>CACATCCCCAGGAGCATAGATAGCGATGTGACTGGGGTAAGCGAAGGCAGCCA<br>ACGCAGCAGCAGCGTGAAAGGCGTCAGGAGTTTTTGAGTTGCCGTGGAAACTT<br>TCAGCCCATCCCAGGATGGGCTTTTTTctccaccaacaatctaga | |
| pP$_{L}$-GlnZ1 G149C | aTCGTATATTAAAAATCCGACAAATTTCGCGTTGCTGCAAGGCAGCAACTGAG<br>CACATCCCCAGGAGCATAGATAGCGATGTGACTGGGGTAAGCGAAGGCAGCCA<br>ACGCAGCAGCAGCGTGAAAGGCGTCAGGAGTTTTTGAGTTGCCCTGGAAACTT<br>TCAGCCCATCCCAGGATGGGCTTTTTTctccaccaacaatctaga | G149C |
| pP$_{L}$-GlnZ1 G146C | aTCGTATATTAAAAATCCGACAAATTTCGCGTTGCTGCAAGGCAGCAACTGAG<br>CACATCCCCAGGAGCATAGATAGCGATGTGACTGGGGTAAGCGAAGGCAGCCA<br>ACGCAGCAGCAGCGTGAAAGGCGTCAGGAGTTTTTGAGTTCCCGTGGAAACTT<br>TCAGCCCATCCCAGGATGGGCTTTTTTctccaccaacaatctaga | G146C |
| pP$_{L}$-GlnZ1 G141U | aTCGTATATTAAAAATCCGACAAATTTCGCGTTGCTGCAAGGCAGCAACTGAG<br>CACATCCCCAGGAGCATAGATAGCGATGTGACTGGGGTAAGCGAAGGCAGCCA<br>ACGCAGCAGCAGCGTGAAAGGCGTCAGGAGTTTTTAGTTGCCGTGGAAACTTT<br>CAGCCCATCCCAGGATGGGCTTTTTTctccaccaacaatctaga | G141U |
| pP$_{L}$-GlnZ1 G141U/G146C | aTCGTATATTAAAAATCCGACAAATTTCGCGTTGCTGCAAGGCAGCAACTGAG<br>CACATCCCCAGGAGCATAGATAGCGATGTGACTGGGGTAAGCGAAGGCAGCCA<br>ACGCAGCAGCAGCGTGAAAGGCGTCAGGAGTTTTTAGTTCCCGTGGAAACTTT<br>CAGCCCATCCCAGGATGGGCTTTTTTctccaccaacaatctaga | G141U/G146C |
| pP$_{L}$-GlnZ2 | aGTTTTTGAGTTGCCGTGGAAACTTTCAGCCCATCCCAGGATGGGCTTTTTTc<br>tccaccaacaatctaga | |
| pP$_{L}$-GlnZ$_{K12}$ | aGTGTTTTAGTTGCCGTGGAAACTTTTCGCCTGTCTCTGGCAGGCCTGGGATC<br>GGTGGCAAGCACATCACGCCGGATGCGACGCAAATGCGTCTTATCCGGCCTAC<br>ACGGTGATGATGTGGTAGGCCGGAGCAGGTGAGTCGCTCTCCAACGTGAAGTT<br>TGTCAGCTATCTGTAGCCCATCTCTGCATGGGCTTTTTTctccgtcaattcta<br>ga | |
| pP$_{L}$-GlnZ$_{K12}$ G15C | aGTGTTTTAGTTGCCCTGGAAACTTTTCGCCTGTCTCTGGCAGGCCTGGGATC<br>GGTGGCAAGCACATCACGCCGGATGCGACGCAAATGCGTCTTATCCGGCCTAC<br>ACGGTGATGATGTGGTAGGCCGGAGCAGGTGAGTCGCTCTCCAACGTGAAGTT<br>TGTCAGCTATCTGTAGCCCATCTCTGCATGGGCTTTTTTctccgtcaattcta<br>ga | G15C |
| pP$_{L}$-GlnZ$_{K12}$ G12C | aGTGTTTTAGTTCCCGTGGAAACTTTTCGCCTGTCTCTGGCAGGCCTGGGATC<br>GGTGGCAAGCACATCACGCCGGATGCGACGCAAATGCGTCTTATCCGGCCTAC<br>ACGGTGATGATGTGGTAGGCCGGAGCAGGTGAGTCGCTCTCCAACGTGAAGTT<br>TGTCAGCTATCTGTAGCCCATCTCTGCATGGGCTTTTTTctccgtcaattcta<br>ga | G12C |
| pP$_{L}$-GlnZ$_{O157}$ | aTAGTTGAAGTTGTACTACCCGGCGCAACAACGCCGGGATTTAGTTGCCGTGG<br>AAACTTTCAGCCCATCTCTGCATGGGCTTTTTTctccgtcaattctaga | |
| pP$_{L}$-GlnZ$_{O157}$ G49C | aTAGTTGAAGTTGTACTACCCGGCGCAACAACGCCGGGATTTAGTTGCCCTGG<br>AAACTTTCAGCCCATCTCTGCATGGGCTTTTTTctccgtcaattctaga | G49C |
| pP$_{L}$-GlnZ$_{O157}$ G46C | aTAGTTGAAGTTGTACTACCCGGCGCAACAACGCCGGGATTTAGTTCCCGTGG<br>AAACTTTCAGCCCATCTCTGCATGGGCTTTTTTctccgtcaattctaga | G46C |
| pP$_{L}$-GlnZ$_{O157}$ U41G | aTAGTTGAAGTTGTACTACCCGGCGCAACAACGCCGGGATTGAGTTGCCGTGG<br>AAACTTTCAGCCCATCTCTGCATGGGCTTTTTTctccgtcaattctaga | U41G |
| pP$_{L}$-GlnZ$_{O111}$ | aTAGTTGAAGTTGTACTACCCGGCGCAACAACGCCGGAGGTTCAAACCAGGCC<br>CATCTGGCGTAATTGTTGCAGCCAGTTTGAACACGGACAGCGCGCAGAACCCG<br>GAGCGTACACTGGTACGTGAGGAGTTCGAGCACTGCCCAGGTTCAAAATGGCA<br>AATAAAATAGCCTGATGGGACTGGTTTTTAGTTGCCGTGGAAATTTCAGCCCA<br>TCTCTGCATGGGCTTTTTTctccgtcaattctaga | |

### Table S6. Details of GFP fusion plasmids.

| Target gene | Vector | Oligonucleotide | Upstream ORF [bp] | Intergenic region [bp] | Downstream ORF [bp] | Insert length [bp] | Translational fusion to N-terminal FLAG [aa] | Translational fusion to C-terminal GFP [aa] |
|---|---|---|---|---|---|---|---|---|
| *sdhC-sucAsal* | pXG-30sf | MMO-0325 x MMO-0326 | 120 | 523 | 30 | 673 | 40 | 10 |
| *sdhC-sucAeco* | pXG-30sf | MMO-0325 x MMO-0326 | 120 | 300 | 30 | 450 | 40 | 10 |
| *deoBDsal* | pXG-30sf | MMO-0529 x MMO-0530 | 30 | 209 | 60 | 299 | 10 | 20 |
| *glnHPsal* | pXG-30sf | MMO-0594 x MMO-0595 | 42 | 143 | 15 | 200 | 14 | 5 |
| *pdhR-aceEsal* | pXG-30sf | MMO-0701 x MMO-0702 | 189 | 159 | 90 | 438 | 63 | 30 |
| *pdhR-aceEeco* | pXG-30sf | MMO-0701 x MMO-0702 | 189 | 160 | 90 | 439 | 63 | 30 |

### Table S7. Inserts of GFP fusion plasmids.

Nucleotide sequences of upstream ORF fused with FLAG, intergenic region, and downstream ORF fused with GFP are indicated in blue, black, and red, respectively. The mutated nucleotides in the GlnZ target region are highlighted in magenta. NsiI and NheI sites used for cloning are highlighted in bold in cyan and green, respectively.

| GFP fusion | Insert |
|---|---|
| *sdhC-sucAsal* | ATGCATGCATTCAGCGTATTCCGCTGTCACAGCATTATGAACTGCGTCAGTGTATGTCCTAAGGGACTGAA<br>CCCGACGCGCGCTATCGGCCATATTAAGTCGATGCTGTTGCAGCGTAGCGCATAAGTAGTTGTAGTGGTTG<br>CCGGATGGCGACGTACAGTCTTATCCGGCCTACAGGCGCAACCCCGTAGGCCTGATAAGCGCAGCGCCATC<br>AGGCAATAGAAGTGCAGGAAACCTCTAAAAACTGCCATATGATTAGTAATAATCGAGATGTTCAGAGCGAG<br>ACAAGGCGCGGTCTGAACGAATCTTCGGGAGCATAGTTAACTATGTGACCGGAGTGAGCGAAGGCCGCAAC<br>GCAGTCGTAGCCTGAACAGCGAAGATTATTAGACAGTTTTTAAAGGTTCCTTAGCGGACGCAGATGACACG<br>ACAGTCCGCAACAAGTGAACCCCGGCACGCATACAGCGTATGCGTGGTTAGTATCCACGGCGAAATACTCG<br>TCATAGTTCACGTTGCATGTGCGTTGGCTGCACTTGCTCACCCCGGTCACTTACTGGATGTAAGCTCCCAG<br>GGATTCGCAAGCTTGCCGCCTTCCTGAAACGTGACCTATTTAGAGTATTAAATAAGCAGAAAAGATGCTTA<br>AGGGATCACGATGCAGAACAGCGCTTTGAAAGCCTGGTTGGCTAGC |
| *sdhC-sucAeco* | ATGCATGCATTCAGCGTATTCCGCTGTCACAGCATCATGAACTGCGTCAGTGTATGTCCGAAGGGGCTGAA<br>CCCGACGCGCGCCATCGGCCATATCAAGTCGATGTTGTTGCAACGTAATGCGTAAACCGTAGGCCTGATAA<br>GACGCGCAAGCGTCGCATCAGGCAACCAGTGCCGGATGCGGCGTGAACGCCTTATCCGGCCTACAAGTCAT<br>TACCCGTAGGCCTGATAAGCGCAGCGCATCAGGCGTAACAAAGAAATGCAGGAAATCTTTAAAAACTGCCC<br>CTGACACTAAGACAGTTTTTAAAGGTTCCTTCGCGAGCCACTACGTAGACAAGAGCTCGCAAGTGAACCCC<br>GGCACGCACATCACTGTGCGTGGTAGTATCCACGGCGAAGTAAGCATAAAAAAGATGCTTAAGGGATCACG<br>ATGCAGAACAGCGCTTTGAAAGCCTGGTTGGCTAGC |
| *deoBDsal* | ATGCATCCGATGGACTACGGTAAAAACATGCTGTGATACCTGTTGTCTTTCGCGCTGTAGATGCGTTGACT<br>TCATTCGCTCACCCCGGTCACATAGTTAATCTATGCTCCCGGGGATTCACGAATTCGTCGCCTTCCTACAA<br>CGCGAAATACTTAGGGTATTTCACTGTTTTGACACACGTTTTGTAGGCCTGATAAGGCGTAGCCGCCATCA<br>GGCAATAACAACAAATTTAAAGGGAACTGAAGATGGCAACTCCACATATTAATGCAGAAATGGGTGATTTC<br>GCTGACGTCGTATTGATGCCG<br>GCTAGC |
| *glnHPsal* | ATGCATAACGAAATCTACAAAAAATGGTTCGGTACAGAACCTAAATAAAACGACCTGATGTTGGCTTGTGG<br>CCCGGTAGTGAGGACGCTCACCGGGCCTACAGCTCTTTCCGTGACGGATAAGACACTTTCCGCCATTTCCG<br>CGCCGGATAGAGCATTTGACTATTTACACCACGGTAACAGGAACGACATATGCAGTTTGACTGGGCTAGC |
| *pdhR-aceEsal* | ATGCATGAGATGCTGCCGCTGGTGAGTACGCATCGCACCCGTATATTTGAAGCCATTATGGCCGGAAAACC<br>AGAAGAAGCGCGTGAAGCGTCACACCGACATCTGGCGTTTATCGAAGAGATTATGCTGGACAGAAGCCGTG<br>AGGAGAGCCGTCGTGAACGCGCTTTACGCCGCCTGGAACAGCGCAAGAATTAGTTTTTTCTGGCAGAACGT<br>TAACCAGAAGATGTTGTGAATAACGCGCAAAAATGTGCGCGGCAACTAAACGCAGAACCTGTCTTATTAAG<br>CTTTCTCACGAAAGTTTAATGGGACAGGTTCCAGATAACTCAACGTATTAGATAGATAAGGAATACCCCCA<br>TGTCAGAACGTTTCCAAAATGACGTGGATCCGATCGAAACTCGCGACTGGCTACAGGCGATCGAATCGGTC<br>ATCCGTGAAGAAGGTGTTGCTAGC |
| *pdhR-aceEeco* | ATGCATGAGATGCTGCCGCTGGTGAGTAGTCACCGCACCCGCATATTTGAAGCGATTATGGCCGGTAAGCC<br>GGAAGAAGCGCGCGAAGCATCGCATCGCCATCTGGCCTTTATCGAAGAAATTTTGCTCGACAGAAGTCGTG<br>AAGAGAGCCGCCGTGAGCGTTCTCTGCGTCGTCTGGAGCAACGAAAGAATTAGTGATTTTTCTGGTAAAAA<br>TTATCCAGAAGATGTTGTAAATCAAGCGCATATAAAAGCGCGGCAACTAAACGTAGAACCTGTCTTATTGA<br>GCTTTCCGGCGAGAGTTCAATGGGACAGGTTCCAGAAAACTCAACGTTATTAGATAGATAAGGAATAACCC<br>ATGTCAGAACGTTTCCCAAATGACGTGGATCCGATCGAAACTCGCGACTGGCTCCAGGCGATCGAATCGGT<br>CATCCGTGAAGAAGGTGTTGCTAGC |
